# High genetic load without purging in a diverse species-at-risk

**DOI:** 10.1101/2022.12.19.521038

**Authors:** Rebecca S. Taylor, Micheline Manseau, Sonesinh Keobouasone, Peng Liu, Gabriela Mastromonaco, Kirsten Solmundson, Allicia Kelly, Nicholas C. Larter, Mary Gamberg, Helen Schwantje, Caeley Thacker, Jean Polfus, Leon Andrew, Dave Hervieux, Deborah Simmons, Paul J. Wilson

## Abstract

High intra-specific genetic diversity is associated with adaptive potential which is key for resilience to global change. However, high variation may also support deleterious alleles through genetic load, unless purged, thereby increasing the risk of inbreeding depression if population sizes decrease rapidly. Purging of deleterious variation has now been demonstrated in some threatened species. However, less is known about the costs of population declines and inbreeding in species with large population sizes and high genetic diversity even though this encompasses many species globally that have or are expected to undergo rapid population declines. Caribou is a species of ecological and cultural significance in North America with a continental-wide distribution supporting extensive phenotypic variation, but with some populations undergoing significant declines resulting in their at-risk status in Canada. We assessed intra-specific genetic variation, adaptive divergence, inbreeding, and genetic load across populations with different demographic histories using an annotated chromosome-scale reference genome and 66 whole genome sequences. We found high genetic diversity and nine phylogenomic lineages across the continent with adaptive diversification of genes, but also high genetic load among lineages. We also found highly divergent levels of inbreeding across individuals including the loss of alleles by drift (genetic erosion) but not purging, likely due to rapid population declines not allowing time for purging of deleterious alleles. As a result, further inbreeding may need to be mitigated through conservation efforts. Our results highlight the ‘double-edged sword’ of genetic diversity that may be representative of other species-at-risk affected by anthropogenic activities.

## INTRODUCTION

lntra-specific diversity is crucial for adaptive potential and resilience of species under environmental changes (Andrello et al., 2022; Carvalho et al., 2017; O’Brien et al., 2022; Hoban et al., 2022;). Therefore, understanding the drivers of intra-specific genetic variation and its interplay with adaptive divergence is essential to understanding how current species respond to environmental variations (Des Roches et al., 2021; Leigh et al., 2021; Yiming et al., 2021). Conversely, there is growing evidence suggesting that a larger genetic load is present in populations with high genetic diversity (Bertorelle et al. 2022; van Oosterhout et al. preprint). Recent research on threatened populations or species with low genetic diversity has demonstrated the purging of deleterious genetic variation due to the maintenance of small population sizes, for example, in the Sumatran rhinoceros (vonSeth et al. 2022), the kākāpō (Dussex et al. 2021), Alpine ibex (Grossen et al. 2020), and Indian tigers (Khan et al. 2021), with some threatened species nevertheless developing inbreeding depression likely due to rapid declines and historical demography, for example, in killer whales (Kardos et al. 2023) and Scandinavian wolves (Smeds and Ellegren, 2022).

Less is known about the costs of population declines and inbreeding in species with large population sizes and high genetic diversity (Fairmount et al. 2023), even though this encompasses many species globally that have not maintained small population sizes but will likely undergo rapid declines and fragmentation into isolated populations due to anthropogenic impacts. Such species with large historical effective populations sizes are expected to have high genetic load (Bertorelle et al. 2022; van Oosterhout et al. preprint) and may decline too rapidly to purge deleterious variation. We investigate these processes in an example of such a species, caribou (*Rangifer tarandus*), a wide-spread and diverse species-at-risk.

Caribou (known as reindeer in Eurasia) is a highly mobile keystone species with a continental-wide distribution, ranging from the high Arctic to the boreal forests, and spanning from the east to the west coast of North America (COSEWIC, 2011; Figure 1). Across its range caribou have a large amount of phenotypic and genetic variation and have been divided into 12 conservation units, known as Designatable Units (DUs), by the Committee on the Status of Endangered Wildlife in Canada (COSEWIC, 2011; Figure 1). Caribou DUs face threats including habitat destruction and climate change (Festa-Bianchet et al., 2011; Vors & Boyce, 2009; Weckworth et al., 2018), with nine DUs currently listed as endangered or threatened, two as special concern, and one that is extinct (COSEWIC, 2011-2017). Globally, in 2015 the species changed from Least Concern to Vulnerable on the IUCN Red List due to the species undergoing a 40% decline over three generations (IUCN Red List).

**Figure 1.**
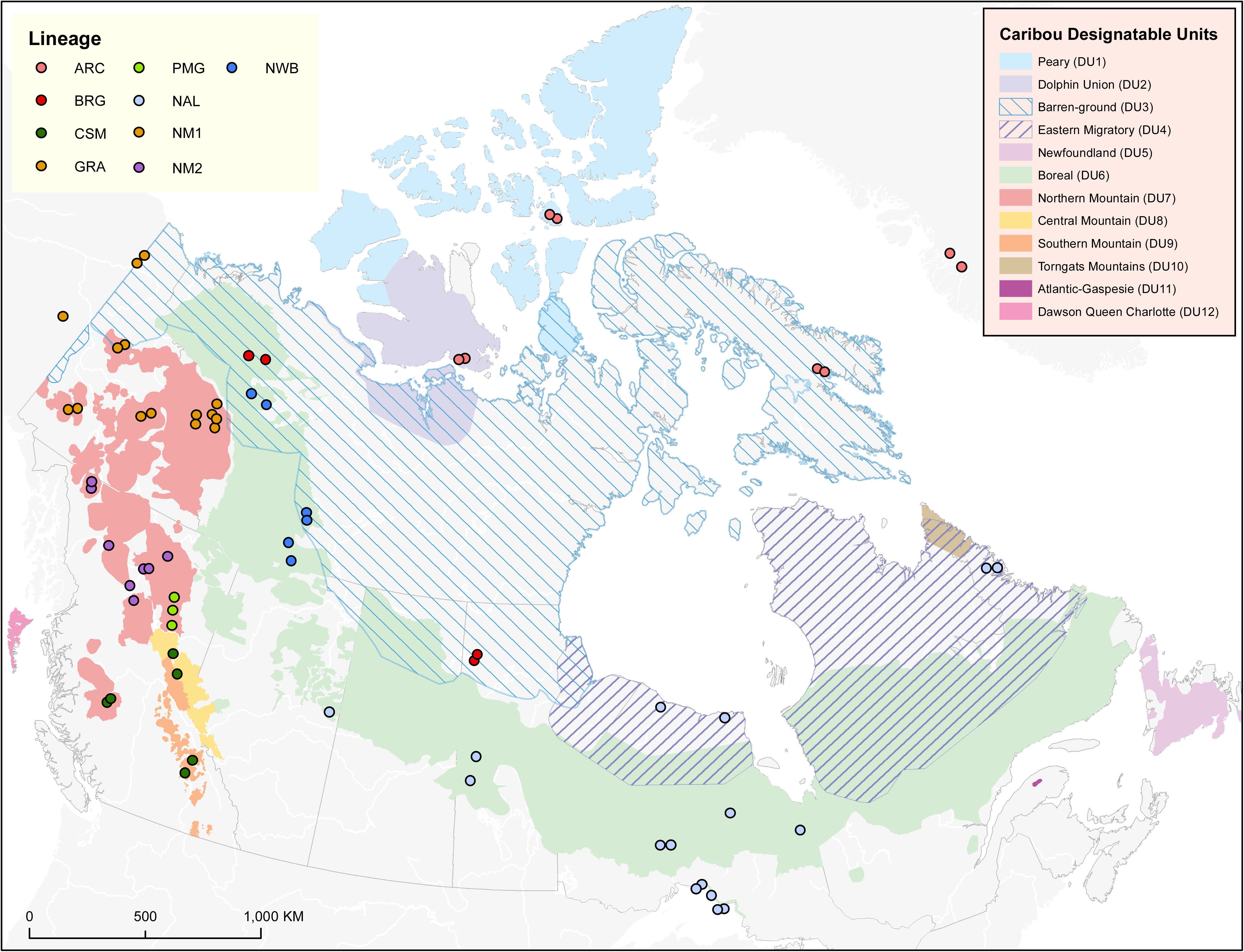
Map of sampled individuals. A map showing the sample location of each of the individuals sequenced. The background shading indicates the Designatable Unit (DU) ranges and the colours of the points indicate which phylogenomic lineage the individual belongs to.

We investigated intra-specific lineage diversity, genetic variation, adaptive diversification, inbreeding extent, and mutational load in caribou across North America and Greenland using whole genome sequencing thus undertaking a comprehensive reconstruction of intra-specific caribou diversity. We first assess intra-specific diversity and the processes that may have led to high genetic variation, as well as adaptive diversification. We then characterize inbreeding and compare genetic load in individuals with different demographic histories (high vs low inbreeding) to understand the potential impact of rapid population declines on genetic diversity as well as on deleterious variation in a genetically and phenotypically diverse species. Since many species have not maintained small population sizes over time but have or are expected to undergo rapid population declines due to anthropogenic impacts including climate change, it becomes imperative to better understand the many facets of genetic diversity and ultimately ensure the long-term resilience of our wild species.

## RESULTS AND DISCUSSION

We assembled a new caribou reference genome with a contig N50 of 32.82 KB, scaffold N50 of 64.42 MB, and an L50 of 14, with 99.5% of the assembly being on 36 scaffolds. As the chromosome number is 70 for the species (34 autosome pairs plus the sex chromosomes; Gripenberg et al. 1986), this likely represents a chromosome-scale assembly. We then used RNA-seq data to perform a high-quality annotation of the genome, which identified the locations of 34,407 protein-coding genes. Using 66 re-sequenced genomes from across North America and Greenland representing eight Designatable Units (DUs) and 33 subpopulations (Figure 1; Table S1 and S2), phylogenomic reconstruction using two different methods was generally consistent and separated caribou into nine major lineages, which were not concordant with DU designations (Figure 2a; Figure S1). We reconstructed the major mitochondrial lineages known from previous studies (Weckworth et al. 2012); the North American lineage (NAL) and the larger and more diverse Beringian-Eurasian lineage (BEL) which contains eight lineages in our results (Figure 2a; Figure S1). The only discordance between our two reconstructions was that the NWB lineage was basal to the PMG lineage instead of a sister group, and that the BRG lineage individuals were within (although on the outside) of the ARC group instead of a sister lineage in the SNP based reconstruction (Figure S2). We show the results based on the more powerful method using full sequence data (Figure 2a), especially given the large differences between BRG and ARC individuals in further analyses. The principal component analysis (PCA) is also concordant with the phylogenomic results separating into three major clusters on PC1: the NAL individuals, NWB, PMG and CSM individuals, and the three major northern mountain lineages (GRA, NM1, and NM2; Figure S3). On PC2, the BRG and ARC lineages separate (Supplementary Figure 3).

**Figure 2.**
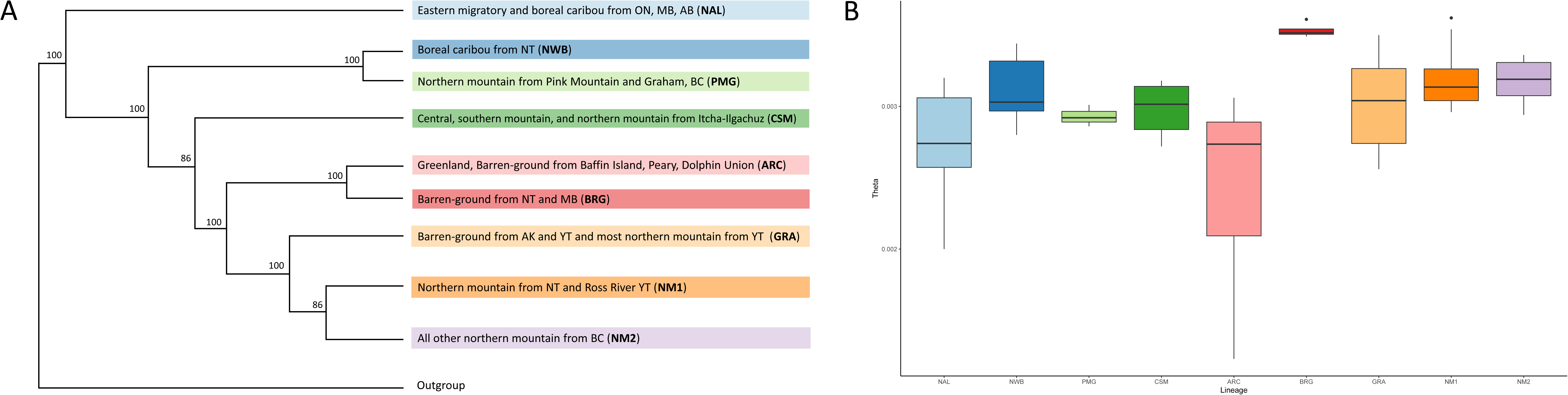
Intra-specific diversity of caribou. **a**, simplified version of the whole genome phylogenomic reconstruction showing which caribou belong to each lineage. **b**, Genetic diversity (θ) for each lineage.

As well as exploring lineage diversity, we calculated individual genetic diversity, θ, an approximation of heterozygosity under the infinite sites model (vonSeth et al. 2022; Haubold et al. 2010; Foote et al. 2021). We found overall high heterozygosity in caribou, although with some variation among individuals (overall mean of 0.0030, range of 0.0012 to 0.0036; Figure 2b; Figure S4). Some individuals from within the NAL and ARC lineages had lower diversity than the others, with the ARC lineage mean θ at 0.0024 and the NAL at 0.0028 compared to the mean of all others at 0.0031 (Figure 2b; Figure S4). When compared to other mammal species where genome-wide heterozygosity has been calculated, our mean is around some of the highest heterozygosity values (see Figure 3 in Morin et al. 2020), demonstrating a high genetic diversity, as well as high phylogenetic lineage diversity, in caribou.

**Figure 3.**
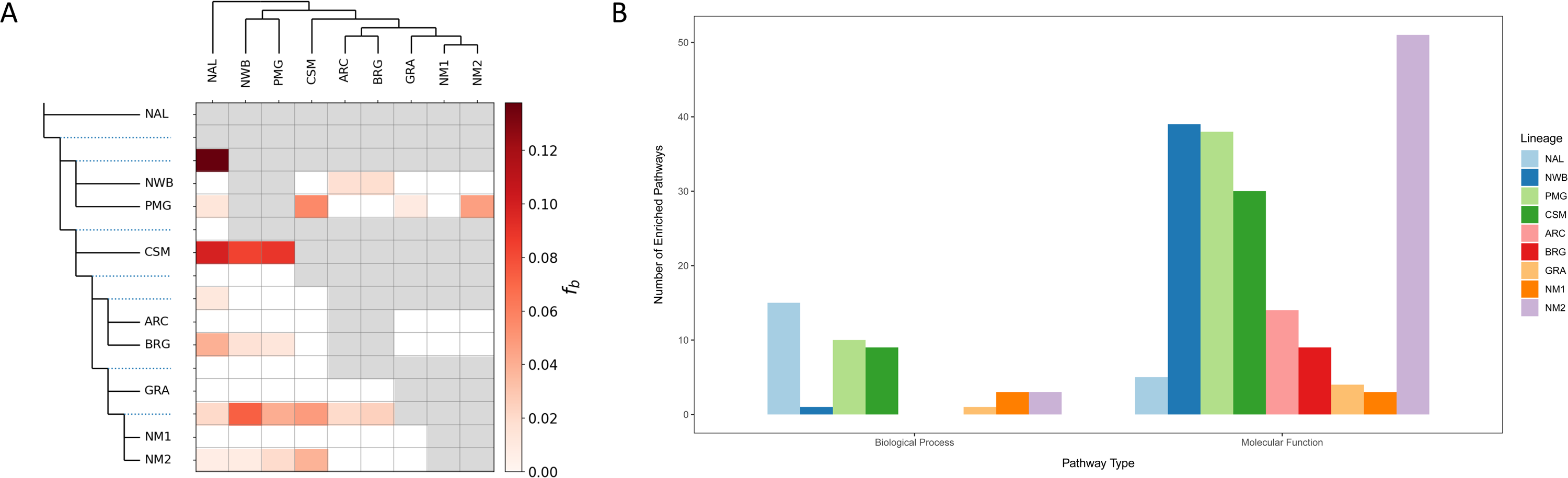
Levels of introgression between lineages and their number of enriched functional pathways. **a,** heatmap showing the f-Branch statistics alongside the phylogeny. The gene flow is from the lineage indicated in the top phylogeny (X-axis) going into the phylogeny represented on the Y-axis. Dotted lines indicate where gene flow is going into an ancestral group on the phylogeny. Greyed out squares indicate tests that could not be made with this statistic, and white squares indicate where no gene flow was detected. **b,** the number of enriched functional pathways within each lineage, under the categories biological processes and molecular function, from their rapidly evolving genes.

To measure introgression among the reconstructed lineages, we used D and f4-ratio tests, which control for incomplete lineage sorting. These statistics gave many significant signatures of introgression between groups (Supplementary Material: Introgression statistics). We then calculated the f-branch statistic, which accounts for many correlated signatures of introgression using the f4-ratio statistics to show when along the phylogeny introgression occurred, and whether the gene flow event was into an ancestral group. The f-branch results indicated widespread introgression between lineages, particularly between NAL and the ancestor of NWB and PMG lineages, and multiple lineages into CSM, the ancestor of NM1 and NM2, NM2, and BRG (Figure 3a). In contrast, some lineages show only a small amount or even a lack of gene flow (with the caveat that sister groups cannot be tested for gene flow using ABBA BABA tests). For example, no lineage shows gene flow into the ARC, GRA, or NM1 caribou lineages (Figure 3a). To explore potential introgression between sister lineages, we visualized ‘admixture graphs’ in SplitsTree. Unsurprisingly these graphs show some phylogenetic uncertainty, putatively due to gene flow, within lineages (Supplementary Figure 5). We also see potential signals of introgression between the NWB, PMG, and CSM lineages, which may help to explain the placement of the PMG individuals next to the NWB lineage in the phylogenetic reconstruction, as well as some potential introgression between the three major northern mountain lineages (GRA, NM1, and NM2).

Altogether our results, reconstructing continent-wide whole genome phylogenomic history for the first time, point towards a high level of intra-specific diversity in caribou, with some strong signals of introgression among many of the lineages. Our results build on previous studies showing high genetic diversity in caribou, for example large numbers of mitochondrial haplotypes (Weckworth et al. 2012; Polfus et al. 2017; Taylor et al. 2021) and high diversity in microsatellite loci (Boulet et al. 2007; McLoughlin et al. 2004; Zittlau et al. 1998). The reasons behind the high diversity and number of intra-specific lineages are likely multi-faceted. The large Beringian refugium, where the individuals from the BEL lineages spent the glacial cycles of the Quaternary, harboured high levels of genetic diversity for some, particularly cold adapted, species such as caribou (Galbreath et al. 2011; Dussex et al. 2020), and is reflected in the large diversity in the BEL vs the NAL caribou (Fig 2a). Post-glacial expansion out of refugia can lead to genetic bottlenecks and low diversity further away from the refugial populations (Roberts and Hamann 2015). However, repeated secondary contact and admixture between glacial lineages can increase genetic diversity (Petit et al. 2003; Alcala and Vuilleumier 2014), in a similar mechanism to the ‘glacial pulse model’ which describes how lineage fusion during glacial cycles can be a source of intra-specific lineage diversity (Maier et al. 2019). Indeed, well-known determinants of diversification during adaptive radiations include high standing variation, gene flow, and habitat to diversify into (Berner and Salzburger 2015), all of which are true for caribou post-glacial recolonizations.

The life history of caribou is also particularly conducive to the formation of high intra-specific variation. Deer species have a high intrinsic rate of increase and dispersal capabilities (Latch et al. 2009). High vagility means caribou were able to undergo range shifts and quickly recolonize habitat once ice sheets recede, or use ice-free corridors (Latch et al. 2009; Dussex et al. 2020; Taylor et al. 2021), facilitating repeated refugial lineage contact potentially increasing genetic diversity as described above.

The same processes leading to high genetic variation will likely also have increased standing adaptive variation in caribou. We used the branch model approach in the codeml module of PAML (Yang 2007) to calculate the ratio of synonymous to non-synonymous mutations (dN/dS ratio) within genes. The program then performs likelihood ratio tests to elucidate whether the ‘focal’ branch has a significantly different ratio from the rest of the phylogenetic tree and is thus putatively a rapidly evolving gene within that branch when compared to the overall phylogeny. We ran each of the nine major linages from our whole genome phylogeny as the focal branch to find those rapidly evolving genes significant to each, thus potentially involved in the adaptive diversification of the lineage.

We found a number of rapidly evolving genes within each lineage (Table S3; Supplementary Material: CodeML). However, contrary to our expectations to find more lineage specific genes in those with lower levels of introgression (as for example in the ARC lineage), we found similar numbers of statistically significant genes after Bonferroni correction for each both overall (X^2^ = 3.058, df = 8, p = 0.931) and for the genes unique to each lineage (X^2^ = 8.142, df = 8, p = 0.420; Table S3). This could be due to the relatively short time scale of the diversification (all within ∼120,000 years; Taylor et al. 2021) limiting the number of genes within each lineage with a high dN/dS ratio. It could also be due to the high level of overall connectivity (introgression) detected among lineages limiting the significant genes resulting from the codeml approach which scans for genes with a strong signature of positive selection when compared to the rest of the phylogeny (i.e. not detecting genes that are important for all or many caribou lineages). There was overlap in the genes that pulled out as significant for each lineage (Supplementary Material: CodeML), which may be indicative of the diversification occurring from a large pool of standing genetic variation, which is known to be a driver of diversification during adaptive radiations (Berner et al. 2015).

We used the codeml approach due to our questions relating to the diversification among caribou lineages, as well as our phylogenomic framework, and have thus characterized some genes putatively involved in the differential adaptation of the caribou lineages (Supplementary Material: CodeML). We performed gene ontology (GO) analyses to assign functional categories to the genes under both ‘biological process’ and ‘molecular function’ and found a number of processes represented in the significant genes such as immune processes, stress responses, carbohydrate binding, amongst many others (Supplementary Material: Gene ontology for all GO terms for each lineage). We then performed enrichment analyses for each lineage, also under both ‘biological process’ and ‘molecular function’, in order to find specific pathways containing multiple genes with signatures of rapid evolution. We found that caribou lineages had different numbers of enriched pathways, with some lineages showing a significantly larger number compared to the others (X^2^ = 112.71, df = 8, p = 2.2e-16; Figure 3b). In particular, the CSM, NWB, and PMG lineages all show high numbers of enriched pathways. These lineages also show some of the highest signatures of introgression, with the CSM lineage showing high introgression from multiple lineages, as well as into the PMG lineage (Figure 3a), and with the admixture graphs showing potential gene flow between the CSM, NWB, and PMG lineages (Figure S5). Similarly, the NM2 lineage also shows a high number of enriched pathways (Figure 3b) and has signatures of introgression from many other lineages (Figure 3a) as well as with the NM1 and GRA lineages (Figure S5). Gene flow may further enhance the variability of functional pathways by exchanging gene variants among lineages creating new combinations, thereby increasing adaptive potential. These combinational pathway changes may well facilitate expression levels and timing prompting adaptation to the range of ecozones inhabited by caribou and the larger *Rangifer* range (Weldenegodguad et al. 2021). It is known that gene flow can facilitate adaptive diversification, as well as inflating standing genetic variation as a whole (Streicher et al. 2014; Berner et al. 2015; Lexer et al. 2016) and may have been a driver of adaptive diversification in these caribou lineages.

Despite their abundance as well as their high phenotypic, lineage, and genetic diversity (Figure 2) with overall high introgression (Figure 3a), and differential adaptive genetic diversity of caribou (Figure 3b; Table S3), some populations have undergone dramatic declines in recent years. For example, the range of boreal caribou in Ontario (NAL lineage) has become disjunct and the populations along the southern edge of the distribution (Lake Superior) have declined to very small numbers of individuals and have already been shown to have elevated signatures of inbreeding (Solmundson et al. 2023). Other populations, for example the Qamanirijuaq barren-ground caribou (BRG lineage) are decreasing but are still in large numbers (∼250,000 individuals for the Qamanirijuaq caribou; COSEWIC, 2016), while some, for example northern mountain caribou from the Redstone (NM1 lineage) and Aishihik (GRA lineage) have remained stable or are increasing (COSEWIC 2014; See Supplementary Material for more detail on what is known about the effective and census population sizes of each of the sampled caribou subpopulations). We measured the effect of demography on signatures of inbreeding using runs of homozygosity (ROH) estimation, and found varied levels of inbreeding across individuals ranging from FROH (proportion of the genome in ROH) of 1% or less in the barren-ground caribou in the BRG and GRA lineages, up to around 56% in caribou from Kangerlussuaq in Greenland (ARC lineage; Figure 4; Figure S6), a number comparable to some of the most endangered species such as southern resident killer whales (Kardos et al. 2023), kākāpō (Dussex et al. 2021) and Indian tigers (Kahn et al. 2021). When we look at the longest ROHs (over 1 million bases long) indicating strong signatures of inbreeding, we see the most in the boreal caribou from the disjunct and most southern part of the range in Ontario (NAL lineage), Itcha-Ilgachuz caribou from British Columbia (CSM lineage), Aishihik caribou from the Yukon (GRA lineage), and Kangerlussuaq caribou from Greenland (ARC lineage; Figure 4; Figure S7). The most inbred caribou are generally from the most northern or southern portions of the distribution where genetic erosion due an extreme environment (north) or anthropogenic disturbance (south) are the strongest (Solmundson et al. 2023) with the exception of the Aishihik caribou which are known to have been introgressed with introduced reindeer (Taylor et al. 2021). For the Itcha-Ilgachuz caribou, declines were seen in the 1900s potentially due to hunting, with the population recovering after the 1970s (Seip and Chichowski 1994; Ministry of Forests, Lands, Natural Resource Operations and Rural Development, 2018), although is once again declining more recently (COSEWIC, 2011-2017). The Kangerlussuaq caribou in Greenland are known to have undergone and strong decline, maintaining low population sizes between 1845 and the 1950s (Cuyler et al. 2002).

**Figure 4.**
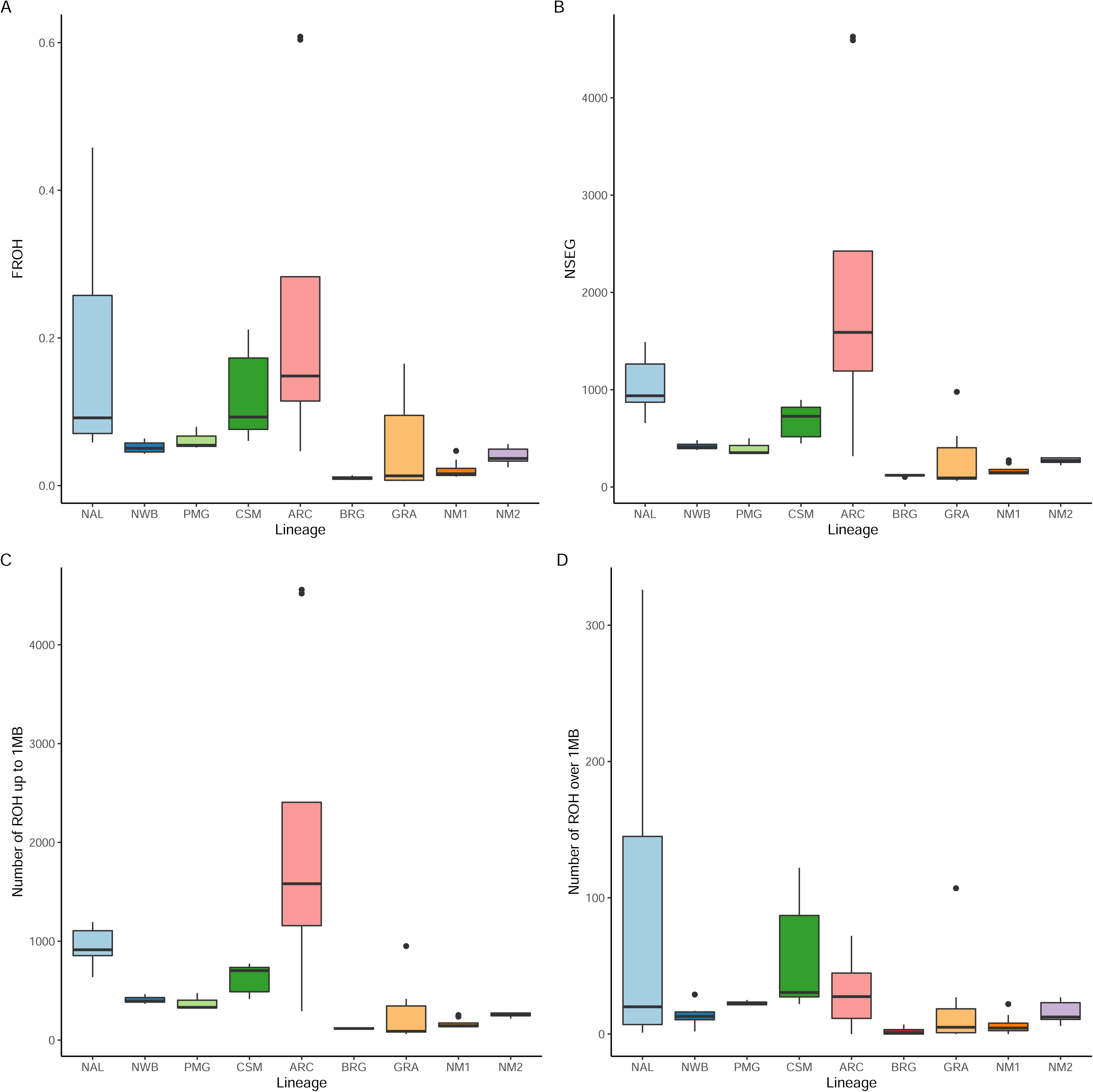
Inbreeding extent in the caribou lineages. **a,** FROH, or the proportion of the genome in runs of homozygosity, **b,** the number of runs of homozygosity in total, **c,** the number of runs of homozygosity up to one million base pairs, and **d,** the number of runs of homozygosity over one million base pairs. The outlier dots for the ARC lineage (**a-c**) are the Kangerlussuaq caribou from Greenland.

Overall, our results demonstrate that different demographic histories have had a dramatic impact on levels of inbreeding and so we investigated genetic load and whether deleterious variation has been purged in the highly inbred compared to non-inbred individuals. Using multispecies comparisons and genomic evolutionary rate profiling (GERP; Davydov et al. 2010), we find fewer derived sites with positive GERP scores and scores over two (at the top end of the score range in our dataset) indicating fewer putatively deleterious variants in the most inbred individuals (Figure 5a and c; Table S4). However, if this was due to purging we would hypothesise the average GERP score of those derived sites to be lower in inbred individuals, but we find no difference in the average score of all positive sites or of the sites with a score over 2 (Figure 5b and c; Table S4) indicating loss of alleles through genetic drift but not purging. Similarly, using our new genome annotation we found fewer derived loss of function (LOF) and high impact alleles in the more inbred individuals, but the pattern is the same for moderate and low impact alleles (Figure 6; Table S4). Our results follow the ‘drift only’ pattern described in Dussex et al. (2023) and thus indicate genomic erosion and loss of overall diversity through drift without preferential purging of deleterious variation. This is in contrast with other species found to have such a high FROH as our most inbred individuals (i.e., 22-42% in disjunct Ontario boreal caribou and 56% in Kangerlussuaq caribou) which have shown to have purged at least some deleterious variation (e.g., Kardos et al. 2023; Dussex et al. 2021; Kahn et al. 2021).

**Figure 5.**
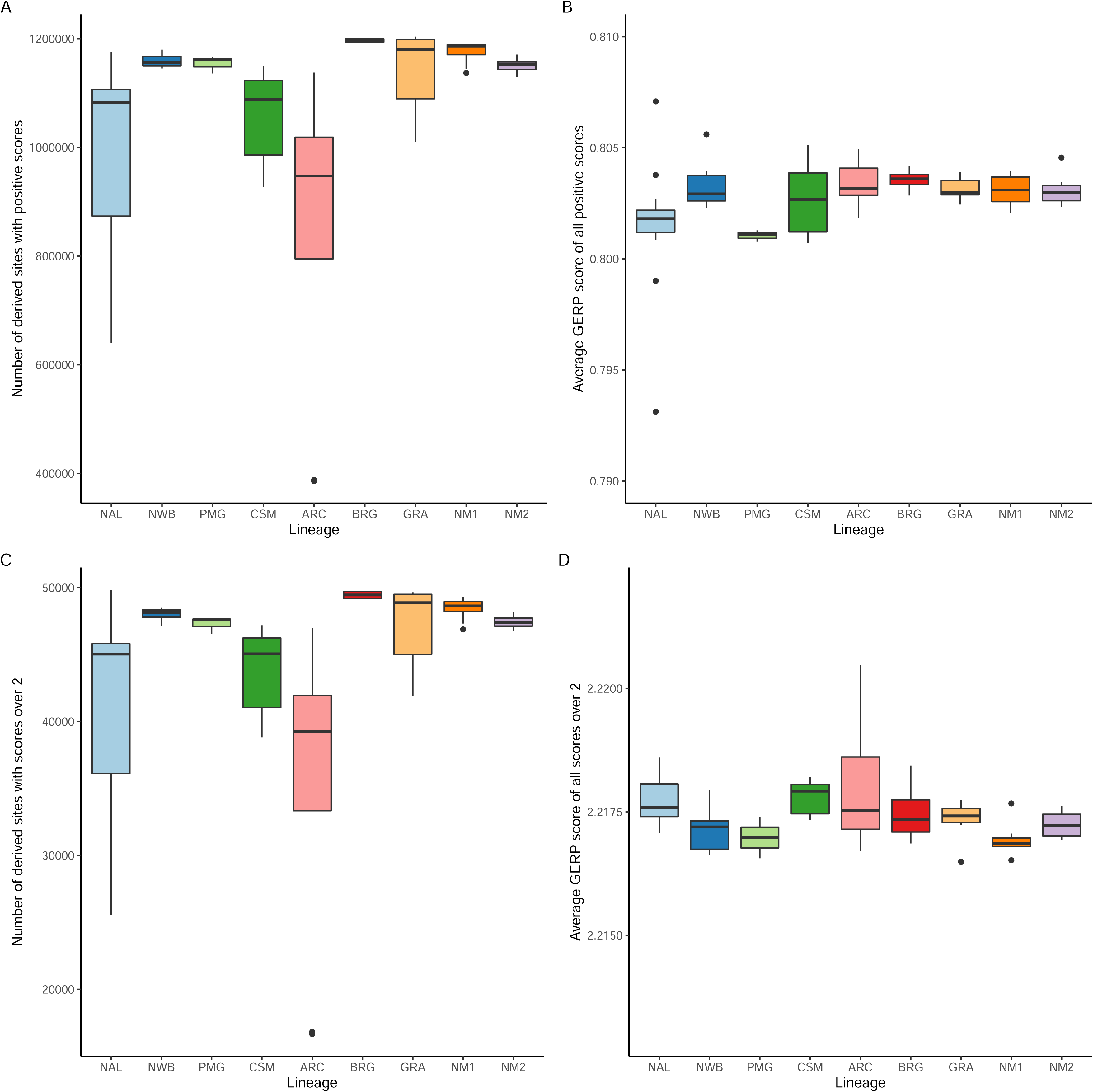
Genetic load as profiled using genomic evolutionary rate profiling (GERP). **a,** the number of derived alleles with a positive GERP score for each lineage and **b,** the average GERP score for the derived alleles with positive scores for each lineage. **c,** the number of derived alleles with a GERP score over two for each lineage and d, the average GERP score for the derived alleles scores over two for each lineage.

**Figure 6.**
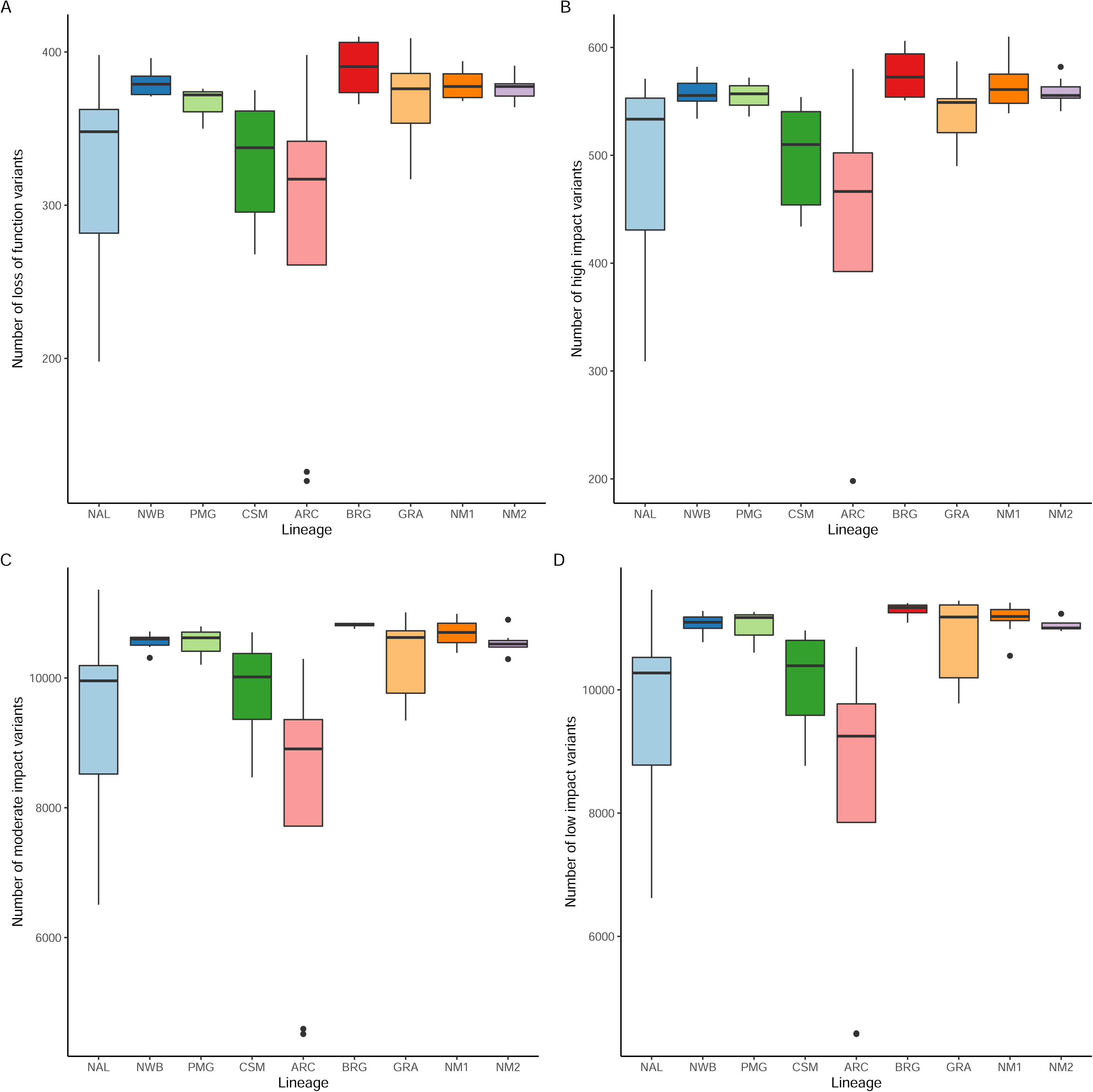
Genetic load as profiled using the genome annotation. The number of derived loss of function, high impact, moderate impact, and low impact alleles in each lineage (**a-d** respectively).

The high impact alleles we uncovered are almost all heterozygous and therefore represent masked load (Bertorelle et al. 2022; van Oosterhout et al. preprint), although we did use strict filtering (e.g., the removal of all sites with any missing data) in order to be conservative and so the numbers are likely underestimates of the ‘true’ load. Even so, we have 11 homozygous high impact alleles representing realized load (Bertorelle et al. 2022; van Oosterhout et al. preprint), eight of which are in the NAL lineage which has high overall inbreeding levels (Table S4), indicating the possibility of these caribou becoming susceptible to inbreeding depression.

It is difficult to compare load across studies due to differences in genome annotations and data filtering, as well as the multi-species alignment used for GERP analysis. However, our data indicate hundreds of high impact alleles present in each individual (between 198 and 610) as well as thousands of moderate impact alleles representing for example, missense mutations (Table S4), representing a large overall genetic load in caribou. This is not surprising given their high historical effective population sizes (Taylor et al. 2021), high phenotypic diversity (COSEWIC), and high genetic and intra-specific lineage diversity and gene flow we reconstructed here.

Preserving the high genetic diversity of caribou, indicated here by the divergent intra-specific lineages and high heterozygosity (Figure 2) and the evidence of differential adaptive variation between lineages (Figure 3; Table S3), may be important for the persistence of caribou and their ability to adapt to environmental changes (Carvalho et al. 2017; Andrello et al. 2022). Given that small populations have not purged deleterious variation, maintaining connectivity between populations and lineages is important as introgression appears to be a driver of genetic variation (Figure 3), allowing movement of adaptive genes (Hanson et al. 2019) and preventing an increase in realized genetic load as has been recommended in other species (e.g., Smeds and Ellegren, 2022). As some of these populations have recently declined to small census sizes, particular attention should be put in assessing the potential impact of inbreeding on current and future trends. Important next steps will include increasing the sample sizes within each subpopulation to enable more detailed population scale analyses, such as recent effective population size reconstructions, a task which will require considerable sequencing effort across such a vast range and will likely need to be undertaken at a regional scale.

### Conservation implications

Much has been published recently in understanding the impacts of low genetic variation and genomic load associated with inbreeding in species with relatively long-term overall low population size (VonSeth et al. 2022; Dussex et al. 2021, Grossen et al. 2020; Khan et al. 2021; Mathur et al. 2023, Kleinman-Ruiz et al. 2022, Xie et al. 2021). However, a different category of at-risk species, such as caribou, sees large effective populations sizes declining thereby manifesting a trade-off of supporting higher potentially adaptive genetic variation, but with the maintenance of non-purged detrimental genetic load with increased probability of expression as such declines occur: the “double edged sword” of higher genetic diversity. We therefore need studies across taxa presenting different demographic histories to enable improved prediction of how a wider variety of species will be affected by population declines.

High genetic load is likely supported in many widespread and diverse species with similar demographic histories to caribou (van Oosterhout et al. preprint) that are perhaps not yet threatened but that have started or will inevitably be impacted by anthropogenic activities such as habitat loss and climate change into the future. As caribou have already begun to be impacted and undergo rapid declines in some areas, the genetic erosion and lack of purging even with high inbreeding levels might foreshadow what will occur in these other taxa.

## METHODS

### Caribou chromosome scale reference genome assembly and annotation

To ensure high quality, contiguous DNA for chromosome-scale reference genome assembly, fibroblast cells were taken from caribou at Toronto Zoo and cultured in T-75 flasks. Firstly, we pre-warmed DMEM1, DMEM3 and trypsin to 37-38 °C, then discarded media and rinsed each T-75 flask with 5 ml of DMEM1. We discarded the media, and added 3 ml of trypsin to each flask and incubate at 38 °C for 2 min. We checked to see that cells had lifted, and then added 9 ml of DMEM3 to each flask, rinsed the flask growing surface to retrieve as many cells as possible and transferred the entire volume to a 15 ml tube, leaving 4 x 15 ml tubes, 2 for each animal. We then centrifuged at 200 x g for 5 min to pellet cells, discarded supernatant and re-suspend pellet in 0.8 ml of PBS. We combined pellets for each individual together in a 2 ml tube, centrifuged in microcentrifuge at 200 x g for 5 min, and discarded the supernatant. The samples were then flash frozen using liquid nitrogen and transferred to a dry shipper and shipped to Dovetail Genomics as we wanted to improve our previous assembly also sequenced by Dovetail Genomics (Taylor et al., 2019), by further scaffolding using Omni-C libraries (Putnam et al., 2016; Yamaguchi et al., 2021). Cells were cultured as above and shipped to Genewiz (Azenta Life Sciences) for RNA sequencing for the annotation. Full details of the sequencing, assembly, and annotation as performed by Dovetail Genomics are available in the Supplementary Material.

### Re-sequenced whole genome sequences

Whole genome sequences of 50 individuals used in this study are available on the National Centre for Biotechnology (NCBI) under BioProject Accession numbers PRJNA634908, PRJNA694662, PRJNA754521, and PRJNA984705 (Table S2; Taylor et al., 2020; Taylor et al., 2021; Taylor et al., 2022). For this study, we sequenced 16 new genomes (Figure 1; Table S2) using the same protocols as before (Taylor et al., 2020; Taylor et al., 2021; details in the Supplementary Material).

All code used to filter and map re-sequenced genomes, as well as for downstream analyses, can be found on GitHub (https://github.com/BeckySTaylor/Phylogenomic_Analyses). Raw reads for all 66 individuals were cleaned using Trimmomatic version 0.38 (Bolger et al., 2014) using a sliding window of 4 base pairs to trim once phred score dropped below 15. We aligned all trimmed reads to the new reference genome, which we first indexed using Bowtie2 version 2.3.0 (Langmead & Salzberg, 2012). We converted the SAM files to BAM files and sorted them using Samtools version 1.5 (Li et al., 2009), and then added read group information using GATK4 (McKenna et al., 2010). Using GATK4, we removed duplicates and used ‘HaplotypeCaller’ to call variants and produce a variant calling format (VCF) file. We used the ‘CombineGVCFs’ function, followed by ‘GenotypeGVCFs’ to produce a VCF file containing all individuals. We did two rounds of filtering on the VCF file using VCFtools version 0.1.16 (Danecek et al., 2011). We removed indels, and any site low-quality genotype calls (minGQ) and low-quality sites (minQ), with scores below 20, as well as any site with a depth of less than five or more than double the mean depth of all genomes, filtering to remove sites with a depth of more than 55. For the second round of filtering, we made two VCF files; one made using a more ‘stringent’ filter to remove all missing data, and a ‘less stringent’ filter to removed sites with more than 5% missing data, resulting in 17,595,673 and 41,321,354 SNPs respectively.

We also downloaded the raw reads for five Fennoscandian Wild Tundra reindeer genomes to use as outgroups for phylogenomic analyses (ID numbers NMBU 38-42 from Weldenegodguad et al., 2020, European Nucleotide Archive accession PRJEB37216). We mapped and filtered the reads as above, as well as producing a VCF file containing these and the 66 caribou genomes. We filtered the VCF file in VCFtools as above, this time removing sites with a depth over 48 (double the mean of this data set) and removing sites with more than 5% missing data, we had 16,119,954 SNPs.

We chose downstream analyses, outlined below, which are appropriate for our sampling across the very large caribou range, which included one or two samples from each of 33 different caribou subpopulations, representing eight DUs (See supplementary materials and Table S1 for more detail on our sampled populations and what is known about their effective and census population sizes). The majority of our analyses are thus those which do not rely on grouping samples together and give individual metrics, with the exception of the CodeML and introgression statistics where we are specifically interested in metrics at a phylogenomic scale. Future work will aim to increase sample size within each subpopulation to enable analyses such as recent Ne reconstruction which need more than 1- 2 samples to run and cannot be done grouping samples with genetic differentiation between them (e.g., using GONE or StairwayPlot2), as well as other analyses requiring the site frequency spectrum. Given the extremely large range of caribou and high number of subpopulations, this will require a huge sequencing effort and likely need to be done at a more regional scale. Many of the results we present here are, however, plotted grouped by lineage for clarity, but all statistics for each individual are given in the supplementary materials.

### Whole genome phylogenomic reconstruction

For the phylogenomic reconstruction, we used IQtree version 1.6.12 (Nguyenet al., 2015). First, we made a consensus fasta file for each individual from the VCF file which included the reindeer using the ‘consensus’ command in BCFtools. We found that running IQtree on the full genome sequences required too much computational power, so we split the genome into seven sections of close to 300 million base pairs, made a phylogeny with each, and then made a consensus tree as follows. Firstly, we used the ‘CSplit’ command to split each individual fasta file into one file per scaffold, retaining the files for scaffolds 1-36 (which contains ∼99% of the reference genome, see results). We concatenated all caribou individuals together for each scaffold, and ran each scaffold from 1-36 in Model Finder in IQtree. Using the Bayesian Information Criterion, Model Finder gave the model TVM+F+I+G4 for all scaffolds apart from 33 where it selected GTR+F+I+G4. For all scaffolds the scores for these two models were close, and for scaffold 33 the likelihood score for the two models was similar (111,917,190.538 and 111,917,210.388), and so when concatenating the scaffolds, we used TVM+F+I+G4 for the full phylogenomic run.

To run the phylogenomic analysis, we then concatenated scaffolds together for each individual into seven fasta files of roughly 300 million base pairs (scaffolds 1-3, 4-7, 8-11, 12-16, 17-21, 22-27, 28- 35), excluding scaffold 36 which is putatively part of the X chromosome based on the presence of known X chromosome genes on that scaffold (Galloway et al., 1996; Liu et al., 2019). We then reformatted each so that the sequence was on one line using ‘awk’ and ‘grep’ commands, and then concatenated all individuals together into one file, including the reindeer, for each of the seven sections so we had one fasta file with all individuals for each of the ∼300 million base pair regions. We then ran IQtree using 100,000 bootstraps to obtain branch supports (-bb command) (Hoang et al., 2018) to produce the phylogenies. We then made a consensus phylogeny from the seven using the IQtree ‘-con’ command.

We also reconstructed an unrooted phylogeny following the protocol from von Seth et al. (2022). We used ngsDist (Vieira et al. 2015) to estimate a genetic distance matrix with 1000 bootstrap replicates from the genotypes allowing no missing data. We then used FASTME v2.1.6.2 (Lefort et al. 2015) to reconstruct the phylogeny, adding bootstrap support to the nodes using RAxML-ng v1.0.1 (Kozlov et al. 2019).

### Principal component analysis, genetic diversity, and introgression measurements

We used Plink version 1.9 (Purcell et al., 2007) to convert the VCF file with the 66 caribou and no missing data into a BED file. We then pruned the dataset to remove sites with a correlation co-efficient of 0.1 or above in sliding windows of 50 SNPs, leaving 3,916,295 putatively unlinked SNPs, and then ran a PCA also in plink, and plotted in R studio version 1.2.5041.

To estimate individual genetic diversity, we used mlRho v2.9 (Haubold et al. 2010) to calculate heterozygosity for each individual from the bam files. mlRho calculates θ, an estimator of the population mutation rate which approximates heterozygosity under the infinite sites model (vonSeth et al. 2022; Haubold et al. 2010; Foote et al. 2021). The files were first filtered using Samtools to remove bases with a mapping quality below 30, sites with a base quality below 30, and with a depth over 10X the average for the dataset.

We measured introgression using ABBA BABA tests to control for incomplete lineage sorting. We used Dsuite version 0.5 (Malinsky et al., 2021) to run the ‘Dtrios’ function to calculate D and f4-ratio statistics, grouping our individuals by the lineages uncovered in our phylogenomic analysis (see results), using the phylogeny as input using the ‘-t’ command. When groups share branches on a phylogeny, many elevated D and f4-ratio statistics can occur, however these correlated statistics can be informative to uncover the relative time of the gene flow events across the phylogeny and to discover whether the gene flow occurred on internal branches by using the f-branch statistic (Malinsky et al., 2021). We calculated the f-branch statistics using the output from Dtrios, and then plotted alongside the phylogeny using the ‘dtools.py’ script included with DSuite, setting the p-value to 0.05.

As these statistics are unable to measure gene flow between sister groups, we used SplitsTree (Hudson and Bryant, 2006) to visualize the phylogenetic network as an ‘admixture graph’, using the seven files SplitsTree output by IQtree (one for each of the 300 million base pair phylogenomic analyses).

### Rapidly evolving genes and gene ontology analysis

We used GWideCodeML (Macías et al., 2020), a python package to run the codeml function of PAML (Yang, 2007) in a computationally efficient way using genome-wide data. We used our annotation file to extract all genes from the genomes of our individuals to use in GWideCodeML. To do this, we made a consensus fasta file for each individual in the VCF file with our 66 caribou filtering to remove sites with more than 5% missing data, as described above. We then used the ‘-x’ function in Gffread version 0.12.3 (Pertea & Pertea, 2020) which pulls out the coding sequence for each gene as indicated in the annotation file and splices them together (to remove introns), to create one fasta file per individual with all genes. We reformatted the files so each gene sequence is on one line using ‘awk’ commands, and then renamed the header line of each gene to include the ID of each individual (in addition to the gene ID from the annotation) using ‘sed’ commands. We used the ‘CSplit’ command to split the files into one file per gene, and then concatenated the files for each gene – resulting in one file per gene containing the sequence for all 66 individuals.

We first ran the genes using an unrooted version of the tree as required by codeml. We removed the outgroup reindeer and then used the ‘ape’ package in R studio to transform the tree into an unrooted version with a trifurcation at the root, the format needed by codeml. We used the branch model, which uses a Likelihood Ratio Test (LRT) to test whether the genes have a significantly different dN/dS ratio on the focal branch, compared to all other branches of the tree. We tested this for each of the nine major lineages uncovered in our phylogenomic analysis, excluding one individual, the boreal caribou from Alberta, which is a hybrid between the NAL and BEL lineages as indicated by the PCA analysis (Supplementary Figure 3). We then performed a Bonferroni multiple testing correction on the Likelihood ratio results, adjusting the significance threshold to account for running the model over nine lineages. We then took those sites where the focal branch was putatively under positive selection (larger dN/dS ratio), and as some genes can be significant in multiple branches, we also calculated how many genes were unique to each branch. We then used a chi-squared test in R studio to determine if there were a significantly different number of positively selected genes across the different lineages.

To assign putative functions to the significant genes (after Bonferroni correction), we used ShinyGo v0.76.2 (Ge et al., 2020) and assigned functions based on the GO Biological Process and GO Molecular Function databases. The enrichment analysis outputs any biological pathways over-represented in the list of genes with signatures of positive selection, and was performed for the significant rapidly evolving genes for each of the nine major lineages separately.

### Runs of homozygosity (ROH) estimation

To estimate the proportion of the genome in ROH we used Plink from the VCF file with no missing data and not LD pruned, and only using scaffolds 1-35 to ensure removing sex chromosomes as above. To test the impact of the key settings (homozyg-snp, homozyg-density, homozyg-gap, homozyg-window-snp, homozyg-window-het, homozyg-het) on the resulting data, we ran 11 different combinations to optimize the runs (Supplementary Material: ROH Plink Settings). Due to our high coverage (over 15X as recommended for this analysis in Plink) and very high SNP density dataset (an average of ∼1 SNP every 125 bp) many of the settings did not affect the results and for those that did we chose a conservative approach (results for all runs available Supplementary Material: ROH Plink Settings) and landed on final settings of homozyg-snp 100, homozyg-density 20, homozyg-gap 1000, homozyg-window-snp 100, homozyg-window-het 1, homozyg-window-missing 5, and homozyg-het 3. Homozyg-kb and homozyg-window-threshold were set using recommendations from Meyermans et al. (2020), so using a homozyg-kb set the same as the scanning window size (100) and using their formula setting homozyg-window-threshold to 0.05.

### Mutational load

We used two approaches to estimate mutational load, one annotation free method and one using our new annotation, to ensure concordance of our results using different approaches and in case of any bias arising from the annotation. For this, we used both genomic evolutionary rate profiling (GERP) analysis (Davydov et al. 2010) and SnpEff (Cingolani et al. 2012) which used our new annotation. For the GERP analysis we largely followed the protocol from von Seth et al. (2022). Firstly, we generated a TimeTree phylogeny (http://www.timetree.org/) of 48 mammal species (Supplementary Figure 8) representing those with available genomes from the even-toed and odd-toed ungulates due to turnover of constrained sites over larger phylogenetic distances (Huber et al. 2020). We downloaded the reference genomes for each of the 48 species and converted to fastq format using BBmap v38.86 (Bushnell et al. 2017), and then aligned to the caribou reference genomes using BWA-MEM (Li, 2013). We converted the resulting alignment files to BAM format and filtered them to remove reads aligning to more than one location as well as supplementary reads, and sorted the resulting file. We then used htsbox (https://github.com/lh3/htsbox) with quality filters (-R -q 30 -Q 30 -l 35 -s 1) to convert into fasta format, and the split each file to make one file per scaffold for the first 36 scaffolds (∼99% of the genome assembly) which were then concatenated together to make one fasta alignment file for each scaffold with all species.

The resulting alignment files were run through a modified version of the gerpcol function in GERP (tar file available here: https://github.com/BeckySTaylor/Phylogenomic_Analyses). Because it can lead to biases (Wootton et al. 2023) the focal species, here caribou, should not be included in the GERP analysis. However, this leads to missing data in the alignment which makes it difficult to interpret the output files which don’t print which site the score pertains to. We modified the code for the gerpcol function to print out the position for each score, as well as the allele for the specified sister species, here the white-tailed deer, which is used as the ancestral allele for each site. This was run using a Ts/Tv ratio of 2.06 as calculated in BCFtools for our caribou dataset. Additionally, as the TimeTree phylogeny outputs the branch lengths in millions of years but gercol requires substitutions per site, we used a tree scaling factor (-s) of 0.0022 reflecting the number of mutations per million years on average per site based on the average mammal mutation rate of 2.2 × 10−9 (Kumar and Subramian, 2002). We then wrote a custom R script (available here: https://github.com/BeckySTaylor/Phylogenomic_Analyses) to automate taking the output and extracting the derived alleles at all sites with positive scores, as well as all sites with a score over 2 (representing the top portion of the possible score range which is a maximum of 2.46 for our dataset and therefore the most highly constrained sites) from our 66 caribou genomes using the VCF file with no missing data.

To get another measure of mutational load, we ran SnpEff using our new annotation and then pulled out SNPs labelled as loss of function (LOF), high impact (which includes the LOF), moderate impact (e.g. missense variants), and low impact (e.g. synonymous variants). We extracted derived SNPs only using the white-tailed deer as an outgroup using the same custom R script as above from the GERP analysis.

### Data and code availability

Whole genome sequences from 50 samples used for this study are available on the National Centre for Biotechnology (NCBI) under BioProject Accession numbers PRJNA634908, PRJNA694662, PRJNA754521, and PRJNA984705. The sequences for the new genomes, and the new reference genome assembly and annotation, will be made available upon acceptance. Bioinformatic code used in this study is available at: https://github.com/BeckySTaylor/Phylogenomic_Analyses

## Supporting information

Plink settings

CodeML

Introgression statistics

Gene Ontology results

## Acknowledgments

We would like to thank the many field collectors and hunters who supplied samples for this work, as well as Bridget Redquest and Austin Thompson for technical support in the laboratory. We would also like to thank Brody Crosby for support with data management, and other members of the Ecogenomics team (https://www.ecogenomicscanada.ca/) for the many insightful conversations. We are also thankful to the Shared Hierarchical Academic Research Computing Network (SHARCNET: www.sharcnet.ca), Compute Canada, and Amazon Cloud Computing for high-performance computing services. Funding from this research was provided by the Genomic Applications Partnership Program of Genome Canada, Environment and Climate Change Canada, the Government of Canada’s Genomics Research and Development Initiative (GRDI), the Species-at-Risk Stewardship Program (SARSP), WCS Garfield Weston Fellowship. We would also like to thank Lukas Keller and two anonymous reviewers for their feedback on the manuscript.

## Author contributions

R.S.T. helped to conceive the study, did the bioinformatics, and wrote the manuscript. M.M. helped to conceive the study, secured funding, and edited the manuscript. S.K. helped with the bioinformatics and edited the manuscript, P.L. wrote scripts for some bioinformatic analyses, and G.M. did the laboratory work to produce the fibroblast cells for the reference genome and annotation. K.S. secured funding and support with bioinformatics, A.K., N.C.L., M.G., H.S., C.T., J.P., L.A., D.H., and D.S. coordinated or collected samples and edited the manuscript. P.J.W. helped to conceive the study, secured funding, and edited the manuscript.

## References

1. Hoban, S. et al. Global genetic diversity status and trends: towards a suite of Essential Biodiversity Variables (EBVs) for genetic composition. Biol. Rev. 97, 1511–1538 (2022).

2. Andrello, M. et al. Evolving spatial conservation prioritization with intraspecific genetic data. TREE 37, 553–564 (2022).

3. Carvalho, S. B. et al. Spatial conservation prioritization of biodiversity spanning the evolutionary continuum. Nat. Ecol. Evol. 1, 0151 (2017).

4. O’Brien, D. et al. Bringing together approaches to reporting on within species genetic diversity. J. Appl. Ecol. 59, 2227–33 (2022).

5. Des Roches, S., Pendleton, L. H., Shapiro, B., & Palkovacs, E. P. Conserving intraspecific variation for nature’s contributions to people. Nat. Ecol. Evol. 5, 574–82 (2021).

6. Leigh, D. M. et al. Opportunities and challenges of macrogenetic studies. Nat. Rev. Genet. 22, 791–807 (2021).

7. Yiming, L. et al. Latitudinal gradients in genetic diversity and natural selection at a highly adaptive gene in terrestrial mammals. Ecography 44, 206–218 (2021).

8. Bertorelle, G. et al. Genetic load: genomic estimates and applications in non-model animals. Nat. Rev. Genet. 23, 492–503 (2022).

9. van Oosterhout, C. et al. Genomic erosion in the assessment of species extinction risk and recovery potential. Preprint at: Genomic erosion in the assessment of species extinction risk and recovery potential | bioRxiv (2022).

10. von Seth, J. et al. Genomic insights into the conservation status of the world’s last remaining Sumatran rhinoceros populations. Nat. Comms. 12, 2393 (2022).

11. Dussex, N. et al. Population genomics of the critically endangered kākāpō. Gell Genomics 1, 100002 (2021).

12. Grossen, C., Guillaume, F., Keller, L. F. & Croll, D. Purging of highly deleterious mutations through severe bottlenecks in Alpine Ibex. Nat. Comms. 11, 1001 (2020).

13. Khan, A. et al. Genomic evidence for inbreeding depression and purging of deleterious genetic variation in Indian tigers. PNAS 118, e2023018118 (2021).

14. Kardos, M. et al. Inbreeding depression explains killer whale population dynamics. Nat. Ecol. Evol. 7, 675–686 (2023).

15. Smeds, L. & Ellegren, H. From high masked to high realized genetic load in inbred Scandinavian wolves. Mol. Ecol. 32, 1567–1580 (2022).

16. Fairmount, A., Rastas, P., Lv, L. & Merila. Inbreeding depression in an outbred stickleback population. Mol. Ecol. (2023).

17. COSEWIC. Designatable units for caribou (Rangifer tarandus) in Canada. Committee on the Status of Endangered Wildlife in Canada (2011).

18. Festa-Bianchet, M., Ray, J. C., Boutin, S., Côté, S. D. & Gunn, A. Conservation of caribou (Rangifer tarandus) in Canada: An uncertain future. Can. J. Zool. 89, 419–434 (2011).

19. Vors, L. S. & Boyce, M. S. Global declines of caribou and reindeer. Global Change Biol. 15, 2626–2633 (2009).

20. Weckworth, B. V., Hebblewhite, M., Mariani, S. & Musiani, M. Lines on a map: Conservation units, meta-population dynamics, and recovery of woodland caribou in Canada. Ecosphere 9, e02323 (2018).

21. COSEWIC. COSEWIC assessment and status reports available Document search - Species at risk registry (canada.ca)

22. Gunn, A. Rangifer tarandus. The IUCN Red List of Threatened Species 2016: e.T29742A22167140 (2016). Accessed on 10 June 2023.

23. Gripenberg, U. & Nieminen, M. The chromosomes of reindeer (Rangifer tarandus). Rangifer 6, 109 (1986).

24. Weckworth, B. V., McDevitt, A., Musiani, M., Hebblewhite, M. & Mariani, S. Reconstruction of caribou evolutionary history in Western North America and its implications for conservation. Mol. Ecol. 21, 3610–3624 (2012).

25. Haubold, B., Pfaffelhuber, P. & Lynch, M. mlRho – a program for estimating the population mutation and recombination rates from shotgun-sequenced diploid genomes. Mol. Ecol. 19, 277–284 (2010).

26. Foote, A. D. et al. Runs of homozygosity in killer whale genomes provide a global record of demographic histories. Mol. Ecol. 30, 6162–6177 (2021).

27. Morin P. A. et al. Reference genome and demographic history of the most endangered marine mammal, the vaquita. Mol. Ecol. Resour. 21, 1008–1020 (2020).

28. Polfus, J. L., Manseau, M., Klütsch, C. F. C., Simmons, D. & Wilson, P. J. Ancient diversification in glacial refugia leads to intraspecific diversity in a Holarctic mammal. J. Biogeog. 44, 386–396 (2017).

29. Taylor. R. S., et al. Population dynamics of caribou shaped by glacial cycles before the last glacial maximum. Mol. Ecol. 30, 6121–6143 (2021).

30. Boulet, M., Couturier, S., Cote, S. D., Otto, R. D. & Bernatchez, L. Integrative use of spatial, genetic, and demographic analyses for investigating genetic connectivity between migratory, montane, and sedentary caribou herds. Mol. Ecol. 20, 4223–40 (2007).

31. McLoughlin, P. D., Paetkau, D., Duda, M. & Boutin, S. Genetic diversity and relatedness of boreal caribou populations in western Canada. Biol. Conserv. 118, 593–598 (2004).

32. Zittlau, K., Coffin, J., Farnell, R., Kuzyk, G. & Strobeck, C. Genetic relationships of three Yukon caribou herds determined by DNA typing. Rangifer 12, 59–62 (1998).

33. Galbreath, K. E., Cook, J. A., Eddingsaas, A. A. & DeChaine, E. G. Diversity and demography in Beringia: Multilocus tests of paleodistribution models reveal the complex history of Arctic ground squirrels. Evolution 65, 1879–1896 (2011).

34. Dussex, N. et al. Moose genomes reveal past glacial demography and the origin of modern lineages. BMC Genomics 21, 854 (2020).

35. Roberts, D. R. & Hamann, A. Glacial refugia and modern genetic diversity of 22 western North American tree species. Proceedings R. Soc. B: Biol. Sci. 282, 20142903 (2015).

36. Petit, R. J. et al. Glacial refugia: hotspots but not melting pots of genetic diversity. Science 300, 1563–1565 (2003).

37. Alcala, N. & Vuilleumier, S. Turnover and accumulation of genetic diversity across large time-scale cycles of isolation and connection of populations. Proceedings R. Soc. B: Biol. Sci. 281, 20141369 (2014).

38. Maier, P. A., Vandergast, A. G., Ostoja, S. M., Aguilar, A. & Bohonak, A. J. Pleistocene glacial cycles drove lineage diversification and fusion in the Yosemite toad (Anaxyrus canorus). Evolution 73, 2476–96 (2019).

39. Berner, D. & Salzburger, W. The genomics of organismal diversification illuminated by adaptive radiations. Trends Genet. 31, 491–499 (2015).

40. Latch, E. K., Heffelfinger, J. R., Fike, J. A. & Rhodes Jr, O. E. Species-wide phylogeography of North American mule deer (Odocoileus hemionus): cryptic glacial refugia and postglacial recolonization. Mol. Ecol. 18, 1730–45 (2009).

41. Yang, Z. PAML 4: phylogenetic analysis by maximum likelihood. Mol. Biol. Evol. 24, 1586–91 (2007).

42. Weldenegodguad, M. et al. Adipose gene expression profiles reveal insights into the adaptation of northern Eurasian semi-domestic reindeer. Comms. Biol. 4, 1170 (2021).

43. Streicher, J. W. et al. Diversification and asymmetrical gene flow across time and space: lineage sorting and hybridization in polytypic barking frogs. Mol. Ecol. 23, 3273–3291 (2014).

44. Lexer, C et al. Gene flow and diversification in a species complex of Alcantarea inselberg bromeliads. Botan. J. Linn. Soc. 181, 505–520 (2016).

45. Solmundson, K. et al. Whole genomes reveal caribou population structure and inbreeding histories. Ecol. Evol. 13, e10278 (2023)

46. Seip, D. R. & Cichowski, D. B. Population Ecology of caribou in British Columbia. Rangifer, 9, 73–80 (1996).

47 Ministry of Forests, Lands, Natural Resource Operations and Rural Development. Itcha-Ilgachuz and Rainbow caribou herd population and habitat information. 54951 (2018).

48. Cuyler, C., Rosing, M., Linnell, J. D. C., Loison, A., Ingerslev, T. & Landa, A. Status of the Kangerlussuaq-Sisimiut caribou population (Rangifer tarandus groenlandicus) in 2000, West Greenland. Greenland Institute of Natural Resources, Technical Report No. 42 (2002).

49. Davydov, E. V. et al. Identifying a high fraction of the human genome to be under selective constraint using GERP++. PLoS Comput. Biol. 6, e1001025 (2010).

50. Dussex, N., Morales, H. E., Grossen, C., Dalén, L. & van Oosterhout, C. Purging and accumulation of genetic load in conservation. TREE (2023).

51. Hanson, J. O., Fuller, R. A. & Rhodes, J. R. Conventional methods for enhancing connectivity in conservation planning do not always maintain gene flow. J. Appl. Ecol. 56, 913–922 (2019).

52. Mathur, S. et al. An evolutionary perspective on genetic load in small isolated populations as informed by whole genome resequencing and forward-time simulations. Evolution 77, 690–704 (2023).

53. Kleinman-Ruiz, D., et al. Purging of deleterious burden in the endangered Iberian lynx. PNAS 119, e2110614119 (2022).

54. Xie, H. et al. Ancient demographics determine the effectiveness of genetic purging in endangered lizards. Mol. Biol. Evol. 39, msab359 (2021).

55. Taylor, R. S. et al. The caribou (Rangifer tarandus) genome. Genes 10, 540 (2019).

56. Putnam, N. H. et al. Chromosome-scale shotgun assembly using an in vitro method for long-range linkage. Genome Res. 26, 342–350 (2016).

57. Yamaguchi, K. et al. Technical considerations in Hi-C scaffolding and evaluation of chromosome-scale genome assemblies. Mol. Ecol. 30, 5923–34 (2021).

58. Taylor, R. S. et al. The role of introgression and ecotypic parallelism in delineating intraspecific conservation units. Mol. Ecol. 29, 2793–2809 (2020).

59. Taylor, R. S. et al. Whole genome sequences from non-invasively collected caribou faecal samples. Conserv. Genet. Resour. 14, 53–68 (2022).

60. Bolger, A. M., Lohse, M. & Usadel, B. trimmomatic: A flexible trimmer for Illumina sequence data. Bioinformatics 30, 2114–2120 (2014).

61. Langmead, B. & Salzberg, S. L. Fast gapped-read alignment with bowtie 2. Nature Methods 9, 357–359 (2012).

62. Li, H., et al. The sequence alignment/map format and samtools. Bioinformatics 25, 2078–2079 (2009).

63. McKenna, A. et al. The genome analysis toolkit: A MapReduce framework for analyzing next-generation DNA sequencing data. Genome Rese. 20, 1297–1303 (2010).

64. Danecek, P. et al. The variant call format and VCFtools. Bioinformatics 27, 2156–2158 (2011).

65. Weldenegodguad, M. et al. Genome sequence and comparative analysis of reindeer (Rangifer tarandus) in northern Eurasia. Sci. Rep. 10, 1–4 (2020).

66. Nguyen, L-T., Schmidt, H. A., von Haeseler, A. & Minh, B. Q. IQ-TREE: A fast and effective stochastic algorithm for estimating maximum likelihood phylogenies. Mol. Biol. Evol. 32, 268–274 (2015).

67. Galloway, S. M. et al. A linkage map of the ovine X chromosome. Genome Res. 6, 667–677 (1996).

68. Liu, R. et al. New insights into mammalian sex chromosome structure and evolution using high-quality sequences from bovine X and Y chromosomes. BMC Genomics 20, 1000 (2019).

69. Hoang, D. T., Chernomor, O., von Haeseler, A., Minh, B. Q. & Vinh, L. S. UFBoot2: Improving the ultrafast bootstrap approximation. Mol. Biol. Evol. 35, 518–522 (2018).

70. Vieira, F. G., Lassalle, F., Korneliussen, T. S. & Fumagalli, M. Improving the estimation of genetic distances from Next-Generation Sequencing data. Biol. J. Linn. Soc. 117, 139–149 (2015).

71. Lefort, V., Desper, R. & Gascuel, O. FastME 2.0: A Comprehensive, Accurate, and Fast Distance-Based Phylogeny Inference Program. Mol. Biol. Evol. 32, 2798–2800 (2015).

72. Kozlov, A. M., Darriba, D., Flouri, T., Morel, B. & Stamatakis, A. RAxML-NG: a fast, scalable and user-friendly tool for maximum likelihood phylogenetic inference. Bioinformatics 35, 4453–4455 (2019).

73. Purcell, S. et al. plink: A tool set for whole-genome association and population-based linkage analyses. Am. J. Human Genet. 81, 559–575 (2007).

74. R Core Team. R: A language and environment for statistical computing. R Foundation for Statistical Computing (2018).

75. Malinsky, M., Matschiner, M. & Svardal, H. dsuite – Fast D statistics and related admixture evidence from VCF files. Mol. Ecol. Resour. 21, 584–595 (2021).

76. Hudson, D. H. & Bryant, D. Application of phylogenetic networks in evolutionary studies. Mol. Biol. Evol. 23, 254–67 (2006).

77. Macías, L. G., Barrio, E. & Toft, C. GWideCodeML: a python package for testing evolutionary hypotheses at the genome-wide level. G3: Genes, Genomes, Genet. 10, 4369-72 (2020).

78. Pertea, G. & Pertea, M. GFF utilities: GffRead and GffCompare. F1000Research, 9 (2020).

79. Ge, S.X., Jung, D. & Yao, R. ShinyGO: a graphical gene-set enrichment tool for animals and plants. Bioinformatics 36, 2628–9 (2020).

80. Meyermans, R., Gorssen, W., Buys, N & Janssens, S. How to study runs of homozygosity using PLINK? A guide for analyzing medium density SNP data in livestock and pet species. BMC Genom. 21, 94.

81. Cingolani, P. et al. A program for annotating and predicting the effects of single nucleotide polymorphisms, SnpEff: SNPs in the genome of Drosophila melanogaster strain w1118; iso-2; iso-3. Fly 6, 80–92 (2012).

82. Huber, C. D., Kim, B. Y. & Lohmueller, K. E. Population genetic models of GERP scores suggest pervasive turnover of constrained sites across mammalian evolution. Plos Genet. 16, e1008827 (2020).

83. Bushnell, B., Rood, J. & Singer, E. BBMerge—Accurate paired shotgun read merging via overlap. PLoS ONE, 12, e0185056 (2017).

84. Li H. (2013) Aligning sequence reads, clone sequences and assembly contigs with BWA-MEM. Preprint at arXiv:1303.3997v2

85. Wootton, E., Robert, C., Taillon, J., Côté, S. D. & Shafer, A. B. A. Genomic health is dependent on long-term population demographic history. Mol. Ecol. 32, 1943– 1954 (2023).

86. Kumar, S. & Subramian, S. Mutation rates in mammalian genomes. PNAS 99, 803–808 (2002).

