## Supplementary tables and figures for "High genetic load without purging in a diverse species-at-risk"

**Supplementary background: caribou census and effective population sizes**

We have sequenced the genomes of caribou representing 33 different subpopulations across North America and Greenland. Reconstruction of historical Ne using pairwise sequentially Markovian coalescent (PSMC) during previous work shows similar trajectories for caribou across the continent, with large increases in Ne starting around 110,000 years ago (Taylor et al. 2020; Taylor et al. 2021), coinciding with the transition to the last glacial period and a period of cooling (Taylor et al. 2021). The large increase is followed by declines in Ne coinciding with periods of dramatic climate changes and rapid warming events starting around 50,000 years ago, a time when North America lost 72% of its large mammalian genera (Taylor et al. 2021). Caribou Ne was therefore lower going into the Last Glacial Maximum and was stable until PSMC can no longer reconstruct Ne, around 10,000 years ago (Taylor et al. 2021). Future work will aim to reconstruct more recent Ne, however methods to do this, such as GONE and StairwayPlot2 (Nadachowska-Brzyska et al. 2021) require a good sample size per location as you cannot group genetically structured individuals such as found within our phylogenetic lineages, and thus this would require a huge genome sequencing effort for caribou to reach ~10 or more genomes per subpopulation/herd.

For many of the subpopulations for which we have genomes, we do have some information on census sizes (Table S1 below) and population trends which we outline here. All of the information we provide here, unless otherwise cited, comes from the COSEWIC status reports (COSEWIC, 2011-2017).

**Peary Caribou**

Peary caribou are a threatened DU sitting within the ARC lineage. The first counts for Peary caribou were made in the 1960s, and there were an estimated 50,000 caribou at this time. In the late 1980’s the population was at an estimated 22,000, but there was a mass die-off reducing the total population to just 5,400 mature individuals by 1996. The most recent estimated census population is 13,200 mature individuals (2015). There are four subpopulations of Peary caribou, and we include two samples from one of those, the Western Queen Elizabeth subpopulation which appears to have been increasing since the 1990s.

**Dolphin Union caribou**

The Dolphin Union caribou are an endangered DU sitting within the ARC lineage from which we sampled two individuals. This herd underwent a strong decline starting in the early 1900s, eventually starting to increase again in the 1970s with migration returning to the mainland. By 1993 up to 7,000 were once again migrating annually across Coronation Gulf and Dease Strait and in 1997 the population was estimated at 34,558 individuals. However, data suggests a decline of around 50-60% over the last 18 years (as of 2017).

**Barren-ground caribou**

The barren-ground caribou are a threatened DU, with 14-15 subpopulations. The overall census size for adult barren-ground caribou is estimated at 800,000 (as of 2016), down from 2 million in the 1990s. Barren-ground caribou are thought to fluctuate in abundance. Most subpopulations are declining, and here we use samples two samples each from four subpopulations: Bluenose-west, Qamanirjuaq (both BRG lineage), Porcupine (in the GRA lineage), and Baffin Island (in the ARC lineage). Bluenose-West have declined by over 80% and Qamanirjuaq by ~40% during the last three generations. Procupine caribou have increased in recent generations, whereas Baffin Island caribou have undergone a dramatic decline of 98% between 1991 and 2014 and there is a large degree of uncertainty in population estimates, and especially low abundance was found on North Baffin.

**Northern mountain caribou**

Northern mountain caribou are a DU listed as special concern, and is estimated to contain a total of 43,000-48,000 mature individuals as of 2014. We find the northern mountain caribou to be split between four phylogenomic lineages; CSM, GRA, NM1, and NM2. There are 45 subpopulations, and we have one to two genome sequences from each of 12 subpopulations: Aishihik, Atlin, Chase, Frog, Graham, Hart River, Itcha-Ilgachuz, Muskwa, Pink Mountain, Spatzizi, Tay, and Tsenaglode. We also have six genome sequences from a 13^th^ subpopulation, the larger Redstone subpopulation. The population trends for sampled populations are largely unknown, apart from Aishihik which is increasing, Atlin and Redstone which are stable, and Itcha-Ilgachuz which is decreasing.

**Central mountain caribou**

Central mountain caribou are an endangered DU and sit within the CSM lineage, estimated at 469 mature individuals as of 2014. This DU has undergone a large decline of at least 64% over the last three generations. There are 10 subpopulations all of which contain fewer than 250 individuals with two now extirpated. All subpopulations have been undergoing long term declines. We have one genome from each of two subpopulations, Kennedy Siding and Quintette.

**Southern mountain**

Southern mountain caribou are listed as endangered and sit within the CSM lineage, estimated at 1,395 mature individuals as of 2014, and have undergone a 45% decline in the last three generations. There are 15 subpopulations all but one of which have been undergoing declines for the last 27 years and contain fewer than 500 individuals, with two subpopulations now extirpated. We have two genomes from one subpopulation, Columbia North.

**Boreal caribou**

The boreal caribou DU, listed as threatened, covers and extremely large range across Canada, however previous genetic evidence, as well as our current results, indicate that boreal caribou belong to two phylogenomic lineages and the phenotype evolved in parallel (Klütsch et al., 2012; Polfus et al. 2017; Taylor et al. 2020). The boreal caribou from the Northwest Territories are from the NWB lineage and the rest sit within the NAL lineage in our results. In total, it was estimated that there are between 24,722 and 30,513 boreal DU individuals as of 2014, and there are currently 51 subpopulations, most of which are in decline. We have six genomes from the Northwest Territories boreal caribou, and 12 genomes from another six of the subpopulations: Coastal, Nipigon, Far North, Kesagami, Naosap, and Cold Lake.

**Eastern migratory caribou**

The eastern migratory DU sits within the NAL lineage, is listed as endangered, and in total was estimated to contain 170,636 mature individuals as of 2017, with an overall decline of 80% recorded over three generations. There are four subpopulations, and we have two genome sequences each from two of them; The eastern migratory caribou from the Southern Hudson Bay subpopulation and the George River subpopulation – a subpopulation which has undergone a particularly dramatic decline of 99% in three generations. It is known that eastern migratory subpopulations have historically fluctuated, however the George River subpopulation, which used the be the largest-sized subpopulation, is now lower than ever recorded and it is unclear if eastern migratory caribou will increase due to new threats.

**Greenland**

We have two genomes from the Kangerlussuaq-Sisimiut caribou population on Greenland. Historically, the herd was in high numbers between 1815 and around 1845, however there was subsequently a huge decline in numbers, remaining low until the 1950s when numbers began to increase steadily again peaking in the 1970s (Cuyler et al. 2002). The herd has been increasing once again in recent years (Cuyler et al. 2002).

**Supplementary Table 1**. Information on census sizes of the sampled subpopulations, giving the population estimates as per COSEWIC status reports (for Canadian subpopulations) in numbers of mature individuals, both for the most recent estimate and the estimate closest to the sampling time.

| **DU** | **Subpopulation** | **Most recent population estimate** | **Year of most recent estimate** | **Year(s) of sampling** | **Closest population estimate to year sampled** | **Year of closest estimate** |
| --- | --- | --- | --- | --- | --- | --- |
| Peary | Western Queen Elizabeth | 7,300 | 2012-2013 | 1993 | Unknown exactly but there was a catastrophic die off in 1994-5 when the Bathurst and adjacent islands crashed  from 3,155 to 542 | 1994-5 |
| Dolphin Union | Dolphin Union | 18,000 | 2015 | 1986 | 34,558 | 1997 |
| Barrenground | Bluenose West | 15,268 | 2015 | 2013 | 20,465 | 2012 |
| Barrenground | Qamanirjuaq | 264,718 | 2014 | 2007/8 | 348,661 | 2008 |
| Barrenground | Porcupine | 197,000 | 2013 | 2001 | 123,000 | 2001 |
| Barrenground | Baffin Island | 4,856 | 2014 | 1995 | 235,000 | 1991 |
| Barrenground | Fortymile | 40,000 | 2022 | 1994 | 22,000 | 1990 |
| Northern mountain | Aishihik | 1,813 | 2009 | 2002 | 889 | 1997 |
| Northern mountain | Atlin | 514-857 | 2007 | 2006 | 514-857 | 2007 |
| Northern mountain | Chase | 404 | 2009 | 2004 | 301 | 2002 |
| Northern mountain | Frog | 199 | 2001 | 2002/3 | 199 | 2001 |
| Northern mountain | Graham | 637 | 2009 | 2021 | 637 | 2009 |
| Northern mountain | Hart River | 1853 | 2006 | 1999/2000 | 1853 | 2006 |
| Northern mountain | Itcha-Ilgachuz | 1220 | 2012 | 2006 | 1547 | 2007 |
| Northern mountain | Muskwa | 828 | 2007 | 2006 | 828 | 2007 |
| Northern mountain | Pink Mountain | 1145 | 1993 | 2004 | 1145 | 1993 |
| Northern mountain | Spatzizi | 2258 | 1994 | 2003 | 2258 | 1994 |
| Northern mountain | Tay | 2907 | 1991 | 2002/5 | 2907 | 1991 |
| Northern mountain | Tsenaglode | 85-340 | 2008 | 2006 | 85-340 | 2008 |
| Northern mountain | Redstone | >7300 | 2012 | 2013/4 and 2019 | >7300 | 2012 |
| Central mountain | Kennedy Siding | 29 | 2014 | 2018 | 29 | 2014 |
| Central mountain | Quintette | 87 | 2014 | 2021 | 87 | 2014 |
| Southern mountain | Columbia North | 157 | 2013 | 2014 | 157 | 2013 |
| Boreal | Northwest Territories | 6,500 | 2012 | 2013/5 and 2019/20 | 6,500 | 2012 |
| Boreal | Coastal | 492 | 2012 | 1999, 2011-2017 | 492 | 2012 |
| Boreal | Nipigon | 300 | 2012 | 2011/12 | 300 | 2012 |
| Boreal | Far North | Unknown | 2012 | 2009 | Unknown | 2012 |
| Boreal | Kesagami | 492 | 2012 | 2009 | 492 | 2012 |
| Boreal | Naosap | 100-200 | 2012 | 2008/9 | 100-200 | 2012 |
| Boreal | Cold Lake | 150 | 2012 | 2014 | 150 | 2012 |
| Eastern migratory | Southern Hudson Bay | 12,479 | 2011 | 1992 | At least 10,798 | 1994 |
| Eastern migratory | George River | 6,704 | 2016 | 2008 | 27,600 | 2012 |
| Greenland | Kangerlussuaq-Sisimiut | 60,469 | 2018 | 2009 | 98,300 | 2010 |

**Supplementary Methods**

**Reference genome sequencing**

For each Dovetail Omni-C library, chromatin was fixed in place with formaldehyde in the nucleus and then extracted. Fixed chromatin was digested with DNAse I, chromatin ends were repaired and ligated to a biotinylated bridge adapter followed by proximity ligation of adapter containing ends. After proximity ligation, crosslinks were reversed and the DNA purified. Purified DNA was treated to remove biotin that was not internal to ligated fragments. Sequencing libraries were generated using NEBNext Ultra enzymes and Illumina-compatible adapters. Biotin-containing fragments were isolated using streptavidin beads before PCR enrichment of each library. The library was sequenced on an Illumina HiSeqX platform to produce ~30x sequence coverage. Then HiRise used MQ>50 reads for scaffolding.

The input de novo assembly and Dovetail OmniC library reads were used as input data for HiRise, a software pipeline designed specifically for using proximity ligation data to scaffold genome assemblies (Putnam et al., 2016). Dovetail OmniC library sequences were aligned to the draft input assembly using bwa (<https://github.com/lh3/bwa>). The separations of Dovetail OmniC read pairs mapped within draft scaffolds were analyzed by HiRise to produce a likelihood model for genomic distance between read pairs, and the model was used to identify and break putative misjoins, to score prospective joins, and make joins above a threshold. We used the stats.sh script in the program BBMap version 38.42 (Bushnell et al., 2017) to calculate the assembly statistics.

**Reference genome annotation**

Cells were cultured as above and shipped to Genewiz (Azenta Life Sciences) for RNA sequencing for the annotation. Total RNA extraction was done using the QIAGEN RNeasy Plus Kit following manufacturer protocols. Total RNA was quantified using Qubit RNA Assay and TapeStation 4200. Prior to library prep, we performed DNase treatment followed by AMPure bead clean up and QIAGEN FastSelect HMR rRNA depletion. Library preparation was done with the NEBNext Ultra II RNA Library Prep Kit following manufacturer protocols. Then these libraries were run on the NovaSeq6000 platform in 2 x 150 bp configuration. The annotation was also performed by Dovetail Genomics. Repeat families found in the genome assemblies of caribou were identified de novo and classified using the software package RepeatModeler (version 2.0.1). RepeatModeler depends on the programs RECON (version 1.08) and RepeatScout (version 1.0.6) for the de novo identification of repeats within the genome. The custom repeat library obtained from RepeatModeler were used to discover, identify and mask the repeats in the assembly file using RepeatMasker (Version 4.1.0). Coding sequences from *Bos taurus*, caribou (www.caribougenome.ca) and reindeer (Li et al., 2017) were used to train the initial ab initio model for caribou using the AUGUSTUS software (version 2.5.5). Six rounds of prediction optimisation were done with the software package provided by AUGUSTUS. The same coding sequences were also used to train a separate ab initio model for caribou using SNAP (version 2006-07-28). RNAseq reads were mapped onto the genome using the STAR aligner software (version 2.7) and intron hints generated with the bam2hints tools within the AUGUSTUS software. MAKER, SNAP and AUGUSTUS (with intron-exon boundary hints provided from RNA-Seq) were then used to predict for genes in the repeat-masked reference genome. To help guide the prediction process, Swiss-Prot peptide sequences from the UniProt database were downloaded and used in conjunction with the protein sequences from *Bos taurus*, caribou (www.caribougenome.ca), and reindeer to generate peptide evidence in the Maker pipeline. Only genes that were predicted by both SNAP and AUGUSTUS softwares were retained in the final gene sets. To help assess the quality of the gene prediction, AED scores were generated for each of the predicted genes as part of the MAKER pipeline. Genes were further characterised for their putative function by performing a BLAST search of the peptide sequences against the UniProt database, and tRNAs were predicted using the software tRNAscan-SE (version 2.05).

**Re-sequenced whole genome sequences**

Samples were extracted using a Qiagen DNAeasy tissue extraction kit following the manufacturer's instructions (Qiagen). Samples were run on a Qubit fluorometer (Thermo Fisher Scientific) using the High Sensitivity Assay Kit and normalized to 20 ng/µl at a final volume of 50 µl. The DNA was shipped to The Centre for Applied Genomics (TCAG) at the Hospital for Sick Children (Toronto, Ontario) for library preparation and sequencing. The samples were sequenced two per lane on an Illumina HiSeq X. The raw reads for the new genomes will be made available on the NCBI upon acceptance.

**References**

1. Taylor. R. S. et al. Population dynamics of caribou shaped by glacial cycles before the last glacial maximum. *Mol. Ecol.* **30**, 6121–6143 (2021).
2. Taylor, R. S. et al. The role of introgression and ecotypic parallelism in delineating intraspecific conservation units. *Mol. Ecol.* **29**, 2793–2809 (2020).
3. Nadachowska-Brzyska, K., Konczal, M., & Babik, W.  Navigating the temporal continuum of effective population size. *Methods in Ecology and Evolution*, **13**, 22–41 (2022).
4. COSEWIC. COSEWIC assessment and status reports available [Document search - Species at risk registry (canada.ca)](https://species-registry.canada.ca/index-en.html#/documents?documentTypeId=18&sortBy=documentTypeSort&sortDirection=asc&pageSize=10&keywords=rangifer)
5. Polfus, J. L., Manseau, M., Klütsch, C. F. C., Simmons, D. & Wilson, P. J. Ancient diversification in glacial refugia leads to intraspecific diversity in a Holarctic mammal. *J. Biogeog.* **44**, 386–396 (2017).
6. Klütsch, C. F. C., Manseau, M., & Wilson, P. J. Phylogeographical analysis of mtDNA data indicates postglacial expansion from multiple glacial refugia in woodland caribou (*Rangifer tarandus caribou*). PLoS One, **7**, e52661 (2012).
7. Cuyler, C., Rosing, M., Linnell, J. D. C., Loison, A., Ingerslev, T. & Landa, A. Status of the Kangerlussuaq-Sisimiut caribou population (*Rangifer tarandus groenlandicus*) in 2000, West Greenland. Greenland Institute of Natural Resources, Technical Report No. 42 (2002).
8. Putnam, N. H. et al. Chromosome-scale shotgun assembly using an in vitro method for long-range linkage. *Genome Res.* **26**, 342–350 (2016).
9. Bushnell, B., Rood, J., & Singer, E. BBMerge—Accurate paired shotgun read merging via overlap. *PLoS ONE,* **12**, e0185056 (2017).
10. Li, Z. et al. Draft genome of the reindeer (*Rangifer tarandus*). *GigaScience,* **6**, 1–5 (2017).

**Supplementary Table 1**. Samples used for whole genome re-sequencing with their ID numbers (PCID), sampling location including the Canadian province (AB = Alberta, BC = British Columbia, MB = Manitoba, NL = Newfoundland and Labrador, NT = Northwest Territories, NU = Nunavut, ON = Ontario, YT = Yukon Territory) or U.S. state (AK = Alaska) , the Designatable Unit to which they belong, the sample type used for DNA extraction, and the average depth in the VCF files used for analyses with no missing data and when allowing 5% missing data shown in brackets.

| **Caribou PCID** | **Sampling location** | **Latitude/longitude** | **Designatable Unit** | **Canadian Sub-population** | **Lineage** | **Sample type** | **Mean depth** |
| --- | --- | --- | --- | --- | --- | --- | --- |
| 15460 | Lower Keel River, NT | 64.725884, -127.051600 | Northern Mountain | Redstone | NM1 | Tissue | 38 (38) |
| 17825 | Deline, NT | 65.496308, -122.825531 | Boreal | Northwest Territories | NWB | Tissue | 37 (37) |
| 17896 | Drum Lake, NT | 63.891177, -126.276909 | Northern Mountain | Redstone | NM1 | Tissue | 36 (36) |
| 20917 | Ft. Severn, ON | 55.802900, -87.779700 | Eastern migratory | Southern Hudson Bay | NAL | Tissue | 35 (34) |
| 21332 | South Brochet Junction, MB | 57.755310, -101.063060 | Barren-ground | Qamanirijuaq | BRG | Tissue | 36 (36) |
| 21350 | Brochet Junction area, MB | 58.010560, -100.865000 | Barren-ground | Qamanirijuaq | BRG | Tissue | 35 (35) |
| 22832 | Hearst, ON | 51.372380, -84.307190 | Boreal | Far North | NAL | Hair | 19 (19) |
| 23507 | Clearwater Fiord, NU | 66.566670, -67.450000 | Barren-ground | Baffin Island | ARC | Tissue | 14 (13) |
| 23508 | Clearwater Fiord, NU | 66.566670, -67.450000 | Barren-ground | Baffin Island | ARC | Tissue | 21 (21) |
| 24476 | Cold Lake, AB | 54.994044, -110,695547 | Boreal | Cold Lake | NAL | Fecal | 14 (14) |
| 27177 | Colville Lake, NT | 67.075993, -124.668217 | Barren-ground | Bluenose West | BRG | Tissue | 38 (38) |
| 27186 | Fort Good Hope, NT | 66.924900, -126.385300 | Barren-ground | Bluenose West | BRG | Tissue | 36 (36) |
| 27601 | Haines Junction, YT | 61.610434, -137.911647 | Northern Mountain | Aishihik | GRA | Tissue | 17 (17) |
| 27602 | Haines Junction, YT | 61.610434, -137.911647 | Northern Mountain | Aishihik | GRA | Tissue | 16 (15) |
| 27673 | Circle, AK | 64.285646, -143.215339 | Barren-ground | NA, Fortymile herd | GRA | Tissue | 17 (17) |
| 27689 | Nain, NL | 56.913347, -61.716601 | Eastern migratory | George River | NAL | Tissue | 36 (36) |
| 27694 | Nain, NL | 56.913347, -61.716601 | Eastern migratory | George River | NAL | Tissue | 36 (36) |
| 27703 | Dawson, YT | 64.613165, -137.538322 | Northern Mountain | Hart River | GRA | Tissue | 16 (15) |
| 27706 | Dawson, YT | 64.613165, -137.538322 | Northern Mountain | Hart River | GRA | Tissue | 17 (16) |
| 27737 | Old Crow, YT | 67.660545, -140.955926 | Barren-ground | Porcupine | GRA | Tissue | 37 (37) |
| 27738 | Old Crow, YT | 67.660545, -140.955926 | Barren-ground | Porcupine | GRA | Tissue | 37 (37) |
| 27772 | Ross River, YT | 62.918412, -132.461189 | Northern Mountain | Tay | NM1 | Tissue | 18 (18) |
| 27773 | Ross River, YT | 62.918412, -132.461189 | Northern Mountain | Tay | NM1 | Tissue | 22 (22) |
| 28320 | Muskwa, BC | 58.665390, -125.28550 | Northern Mountain | Muskwa | NM2 | Tissue | 25 (25) |
| 28327 | Frog, BC | 57.842710, -126.473040 | Northern Mountain | Frog | NM2 | Tissue | 34 (34) |
| 28330 | Pink Mountain, BC | 57.345540, -123.581560 | Northern Mountain | Pink Mountain | PMG | Tissue | 15 (14) |
| 28332 | Spatzizi, BC | 57.040520, -126.824510 | Northern Mountain | Spatzizi | NM2 | Tissue | 22 (22) |
| 28336 | Chase, BC | 56.585820, -126.117780 | Northern Mountain | Chase | NM2 | Tissue | 19 (18) |
| 28337 | Frog, BC | 57.930390, -126.365520 | Northern Mountain | Frog | NM2 | Tissue | 35 (35) |
| 28348 | Tsenaglode, BC | 58.028320, -129.558730 | Northern Mountain | Tsenaglode | NM2 | Tissue | 23 (23) |
| 28395 | Itcha-Ilgachuz, BC | 52.614600, -124.846290 | Northern Mountain | Itcha-Ilgachuz | CSM | Tissue | 32 (32) |
| 28402 | Itcha-Ilgachuz, BC | 52.796100, -124.735300 | Northern Mountain | Itcha-Ilgachuz | CSM | Tissue | 34 (33) |
| 28575 | Atlin, BC | 59.657170, -132.959110 | Northern Mountain | Atlin | NM2 | Tissue | 35 (35) |
| 28580 | Atlin, BC | 59.812640, -133.145770 | Northern Mountain | Atlin | NM2 | Tissue | 33 (33) |
| 28646 | Columbia North, BC | 51.750000, -118.430000 | Southern Mountain | Columbia North | CSM | Tissue | 35 (35) |
| 28649 | Columbia North, BC | 51.750000, -118.430000 | Southern Mountain | Columbia North | CSM | Tissue | 34 (34) |
| 32527 | Cambridge Bay, NT | 69.163000, -105.228000 | Dolphin Union | Dolphin Union | ARC | Tissue | 22 (22) |
| 32529 | Cambridge Bay, NT | 69.163000, -105.228000 | Dolphin Union | Dolphin Union | ARC | Tissue | 18 (17) |
| 34549 | Cornwallis Island, NT | 75.080000, -95.000000 | Peary | Western Queen Elizabeth | ARC | Tissue | 37 (37) |
| 34550 | Cornwallis Island, NT | 75.080000, -95.000000 | Peary | Western Queen Elizabeth | ARC | Tissue | 37 (37) |
| 34590 | Pen Island, ON | 55.024000, -83.537000 | Eastern migratory | Southern Hudson Bay | NAL | Tissue | 35 (35) |
| 35082 | Deline, NT | 65.661384, -124.565284 | Boreal | Northwest Territories | NWB | Tissue | 37 (36) |
| 35324 | The Pas, MB | 53.156200, -100.733400 | Boreal | Naosap | NAL | Tissue | 34 (33) |
| 35326 | Snow Lake, MB | 54.093300, -100.449700 | Boreal | Naosap | NAL | Tissue | 35 (35) |
| 39590 | Neys Area, ON | 48.8019, -86.6660 | Boreal | coastal | NAL | Tissue | 33 (33) |
| 39650 | Michipicoten Island, ON | 47.751900, -85.745700 | Boreal | Coastal | NAL | Tissue | 35 (35) |
| 39651 | Michipicoten Island, ON | 47.752000, -85.838500 | Boreal | Coastal | NAL | Tissue | 37 (37) |
| 39653 | Pukaskwa National Park, ON | 48.326400, -86.182900 | Boreal | Coastal | NAL | Tissue | 37 (37) |
| 39654 | Cochrane, ON | 50.168700, -80.289700 | Boreal | Kesagami | NAL | Tissue | 36 (36) |
| 41660 | Kangerlussuaq, Greenland | 67.247700, -50.289200 | Greenland | NA | ARC | Tissue | 34 (33) |
| 41667 | Kangerlussuaq, Greenland | 67.942700, -50.340300 | Greenland | NA | ARC | Tissue | 30 (29) |
| 45932 | Nipigon, ON | 50.908, -88.34 | Boreal | Nipigon | NAL | Tissue | 16 (15) |
| 45933 | Nipigon, ON | 50.252, -87.83 | Boreal | Nipigon | NAL | Tissue | 18 (18) |
| 45935 | Keel River, NT | 64.210245, -126.514643 | Northern Mountain | Redstone | NM1 | Tissue | 21 (21) |
| 45936 | Keel River, NT | 64.210245, -126.514643 | Northern Mountain | Redstone | NM1 | Tissue | 19 (19) |
| 45967 | Caribou Flats, NT | 63.669889, -127.969174 | Northern Mountain | Redstone | NM1 | Tissue | 16 (16) |
| 45968 | Caribou Flats, NT | 63.669889, -127.969174 | Northern Mountain | Redstone | NM1 | Tissue | 20 (20) |
| 45994 | Slate Islands, ON | 48.667988, -87.025123 | Boreal | Coastal | NAL | Antler | 18 (17) |
| 48050 | South Slave, NT | 62.14911, -116.21731 | Boreal | Northwest Territories | NWB | Blood | 19 (19) |
| 48057 | South Slave, NT | 60.85677, -116.86862 | Boreal | Northwest Territories | NWB | Blood | 16 (16) |
| 48075 | South Slave, NT | 60.218901, -116.222633 | Boreal | Northwest Territories | NWB | Blood | 22 (22) |
| 48094 | South Slave, NT | 61.870213, -115.970767 | Boreal | Northwest Territories | NWB | Blood | 15 (14) |
| 48100 | Quintette, BC | 54.67372, -121.30541 | Central Mountain | Quintette | CSM | Blood | 19 (19) |
| 48105 | Graham, BC | 56.32065, -122.93355 | Northern Mountain | Graham | PMG | Blood | 22 (22) |
| 48111 | Graham, BC | 56.85783, -123.30222 | Northern Mountain | Graham | PMG | Blood | 16 (16) |
| 48112 | Kennedy Siding, BC | 55.3311, -122.0964 | Central Mountain | Kennedy Siding | CSM | Blood | 23 (23) |

**Supplementary Table 2**. The overall number of significant rapidly evolving genes as well as the number of unique genes for each lineage.

| **Lineage** | **Number of genes with a significant dN/dS ratio** | **Number of unique genes with a significant dN/dS ratio** |
| --- | --- | --- |
| NAL | 764 | 245 |
| NWB | 758 | 250 |
| PMG | 779 | 287 |
| CSM | 787 | 270 |
| ARC | 791 | 282 |
| BRG | 777 | 276 |
| GRA | 749 | 271 |
| NM1 | 777 | 265 |
| NM2 | 742 | 278 |

**Supplementary Table 3**. For each individual, the results of genetic load analyses. For genomic evolutionary rate profiling (GERP) the number sites with a positive GERP score, the average of all sites with a positive GERP score, the number sites with a GERP score over two, and the average of all sites with a GERP score over two. For the SnpEff analysis using our annotation, we show the number of heterozygous
(heterozyg.) and homozygous (homozyg.) sites with high impact, loss of function (LOF), moderate impact, and low impact.

| **Caribou PCID** | **Sampling location** | **Number sites positive GERP score** | **Average score sites positive GERP score** | **Number sites GERP score over 2** | **Average score sites GERP score over 2** | **High impact heterozyg.** | **High impact homozyg.** | **LOF heterozyg.** | **LOF homozyg.** | **Moderate impact heterozyg.** | **Moderate impact homozyg.** | **Low impact heterozyg.** | **Low impact homozyg.** |
| --- | --- | --- | --- | --- | --- | --- | --- | --- | --- | --- | --- | --- | --- |
| 15460 | Lower Keel River, NT | 1185400 | 0.803067 | 48659 | 2.21685 | 539 | 0 | 374 | 0 | 10686 | 100 | 11130 | 52 |
| 17825 | Deline, NT | 1154673 | 0.805601 | 48303 | 2.21733 | 550 | 0 | 371 | 0 | 10400 | 78 | 10729 | 45 |
| 17896 | Drum Lake, NT | 1179316 | 0.803129 | 48603 | 2.21706 | 573 | 0 | 371 | 0 | 10533 | 84 | 11135 | 34 |
| 20917 | Ft. Severn, ON | 1125213 | 0.801615 | 46538 | 2.21747 | 552 | 1 | 354 | 1 | 10056 | 70 | 10737 | 40 |
| 21332 | South Brochet Junction, MB | 1199045 | 0.803678 | 49769 | 2.21717 | 555 | 0 | 376 | 0 | 10740 | 96 | 11260 | 44 |
| 21350 | Brochet Junction area, MB | 1200761 | 0.80352 | 49699 | 2.21751 | 590 | 0 | 405 | 0 | 10651 | 106 | 11301 | 62 |
| 22832 | Hearst, ON | 1111100 | 0.802262 | 45998 | 2.2186 | 521 | 0 | 342 | 0 | 10133 | 65 | 10468 | 41 |
| 23507 | Clearwater Fiord, NU | 983973 | 0.802726 | 40561 | 2.2167 | 491 | 0 | 337 | 0 | 9018 | 77 | 9474 | 45 |
| 23508 | Clearwater Fiord, NU | 954051 | 0.803436 | 39642 | 2.21733 | 457 | 0 | 307 | 0 | 8809 | 67 | 9224 | 29 |
| 24476 | Cold Lake, AB | 1157750 | 0.801153 | 47512 | 2.21788 | 570 | 0 | 393 | 0 | 10535 | 79 | 11271 | 46 |
| 27177 | Colville Lake, NT | 1193906 | 0.802852 | 49203 | 2.21844 | 606 | 0 | 410 | 0 | 10744 | 106 | 11355 | 51 |
| 27186 | Fort Good Hope, NT | 1192465 | 0.804158 | 49160 | 2.21686 | 551 | 0 | 366 | 0 | 10727 | 94 | 11049 | 41 |
| 27601 | Haines Junction, YT | 1009904 | 0.802958 | 41880 | 2.21724 | 498 | 0 | 335 | 0 | 9282 | 63 | 9734 | 47 |
| 27602 | Haines Junction, YT | 1026545 | 0.802448 | 42520 | 2.21757 | 490 | 0 | 317 | 0 | 9277 | 74 | 9756 | 38 |
| 27673 | Circle, AK | 1203640 | 0.802985 | 49590 | 2.21732 | 555 | 0 | 380 | 0 | 10912 | 98 | 11397 | 49 |
| 27689 | Nain, NL | 1053122 | 0.803772 | 44349 | 2.21781 | 552 | 0 | 365 | 0 | 9679 | 63 | 10162 | 35 |
| 27694 | Nain, NL | 1087421 | 0.802114 | 45048 | 2.21801 | 552 | 1 | 361 | 1 | 9852 | 58 | 10223 | 44 |
| 27703 | Dawson, YT | 1152140 | 0.803477 | 47521 | 2.21649 | 544 | 0 | 376 | 0 | 10090 | 88 | 10541 | 55 |
| 27706 | Dawson, YT | 1203411 | 0.80281 | 49646 | 2.21757 | 587 | 0 | 409 | 0 | 10529 | 94 | 11324 | 59 |
| 27737 | Old Crow, YT | 1193042 | 0.80389 | 49408 | 2.21742 | 550 | 0 | 392 | 0 | 10624 | 89 | 11317 | 53 |
| 27738 | Old Crow, YT | 1179950 | 0.803558 | 48873 | 2.21774 | 549 | 0 | 372 | 0 | 10639 | 101 | 11127 | 54 |
| 27772 | Ross River, YT | 1143623 | 0.803973 | 47317 | 2.21767 | 563 | 0 | 381 | 0 | 10297 | 89 | 10513 | 39 |
| 27773 | Ross River, YT | 1136923 | 0.802077 | 46878 | 2.21686 | 548 | 1 | 367 | 1 | 10329 | 83 | 10934 | 54 |
| 28320 | Muskwa, BC | 1145016 | 0.803252 | 47178 | 2.217 | 560 | 1 | 377 | 1 | 10214 | 76 | 10933 | 48 |
| 28327 | Frog, BC | 1170576 | 0.803458 | 48195 | 2.21762 | 556 | 0 | 366 | 0 | 10791 | 106 | 10958 | 46 |
| 28330 | Pink Mountain, BC | 1165729 | 0.801282 | 47639 | 2.21656 | 572 | 0 | 372 | 0 | 10700 | 93 | 11123 | 49 |
| 28332 | Spatzizi, BC | 1159383 | 0.802633 | 47926 | 2.21745 | 581 | 1 | 390 | 1 | 10527 | 88 | 11187 | 47 |
| 28336 | Chase, BC | 1153050 | 0.802572 | 47657 | 2.21694 | 555 | 0 | 377 | 0 | 10466 | 96 | 10959 | 44 |
| 28337 | Frog, BC | 1130012 | 0.803156 | 46775 | 2.21708 | 554 | 0 | 364 | 0 | 10400 | 77 | 11046 | 42 |
| 28348 | Tsenaglode, BC | 1156765 | 0.802339 | 47494 | 2.21746 | 571 | 0 | 383 | 0 | 10391 | 91 | 11022 | 60 |
| 28395 | Itcha-Ilgachuz, BC | 926990 | 0.805108 | 38822 | 2.21733 | 434 | 0 | 268 | 0 | 8399 | 71 | 8720 | 48 |
| 28402 | Itcha-Ilgachuz, BC | 952321 | 0.803993 | 39757 | 2.21798 | 440 | 0 | 285 | 0 | 9086 | 77 | 9279 | 49 |
| 28575 | Atlin, BC | 1151580 | 0.802816 | 47290 | 2.21702 | 541 | 0 | 373 | 0 | 10378 | 92 | 10906 | 48 |
| 28580 | Atlin, BC | 1138200 | 0.804557 | 46981 | 2.21738 | 550 | 0 | 378 | 0 | 10477 | 86 | 10938 | 56 |
| 28646 | Columbia North, BC | 1089732 | 0.803487 | 45170 | 2.21733 | 554 | 0 | 375 | 0 | 9880 | 81 | 10322 | 42 |
| 28649 | Columbia North, BC | 1087346 | 0.801841 | 44938 | 2.21786 | 496 | 0 | 327 | 0 | 9985 | 86 | 10372 | 46 |
| 32527 | Cambridge Bay, NT | 1137925 | 0.804381 | 47005 | 2.21774 | 580 | 0 | 398 | 0 | 10197 | 96 | 10651 | 43 |
| 32529 | Cambridge Bay, NT | 1122826 | 0.802899 | 46132 | 2.21705 | 536 | 0 | 356 | 0 | 10073 | 76 | 10475 | 57 |
| 34549 | Cornwallis Island, NT | 930451 | 0.803978 | 38891 | 2.21718 | 469 | 0 | 327 | 0 | 8858 | 79 | 9206 | 38 |
| 34550 | Cornwallis Island, NT | 940499 | 0.802933 | 38836 | 2.21854 | 464 | 0 | 306 | 0 | 8673 | 85 | 8950 | 39 |
| 34590 | Pen Island, ON | 1102321 | 0.801242 | 45607 | 2.21707 | 555 | 1 | 359 | 0 | 10118 | 70 | 10424 | 33 |
| 35082 | Deline, NT | 1148570 | 0.803947 | 48006 | 2.21662 | 551 | 0 | 376 | 0 | 10500 | 85 | 11057 | 54 |
| 35324 | The Pas, MB | 1083624 | 0.802685 | 45060 | 2.21756 | 539 | 1 | 363 | 1 | 9786 | 62 | 10098 | 27 |
| 35326 | Snow Lake, MB | 839790 | 0.785849 | 35069 | 2.21843 | 571 | 0 | 398 | 0 | 11359 | 0 | 11624 | 0 |
| 39590 | Ignace, ON | 862009 | 0.801806 | 36089 | 2.2172 | 427 | 0 | 282 | 0 | 7928 | 58 | 8104 | 29 |
| 39650 | Michipicoten Island, ON | 884963 | 0.802063 | 36154 | 2.21759 | 397 | 0 | 245 | 0 | 7989 | 58 | 8172 | 28 |
| 39651 | Michipicoten Island, ON | 639629 | 0.799008 | 25537 | 2.21834 | 431 | 1 | 280 | 1 | 8589 | 68 | 8856 | 38 |
| 39653 | Pukaskwa National Park, ON | 1067549 | 0.79312 | 44166 | 2.21724 | 308 | 1 | 198 | 0 | 6463 | 44 | 6593 | 29 |
| 39654 | Cochrane, ON | 386086 | 0.801285 | 16822 | 2.21838 | 496 | 0 | 327 | 0 | 9982 | 66 | 10242 | 38 |
| 41660 | Kangerlussuaq, Greenland | 387909 | 0.804953 | 16655 | 2.22048 | 198 | 0 | 120 | 0 | 4470 | 43 | 4403 | 24 |
| 41667 | Kangerlussuaq, Greenland | 1175244 | 0.801836 | 49841 | 2.21883 | 198 | 0 | 126 | 0 | 4547 | 46 | 4386 | 29 |
| 45932 | Nipigon, ON | 1082213 | 0.807086 | 45039 | 2.21734 | 527 | 0 | 352 | 0 | 10153 | 70 | 10541 | 34 |
| 45933 | Nipigon, ON | 1187193 | 0.800865 | 48889 | 2.21812 | 540 | 1 | 343 | 1 | 9940 | 62 | 10339 | 37 |
| 45935 | Keel River, NT | 1189803 | 0.803605 | 49125 | 2.2168 | 582 | 0 | 394 | 0 | 10486 | 102 | 11348 | 54 |
| 45936 | Keel River, NT | 1190111 | 0.802629 | 49288 | 2.21694 | 546 | 0 | 368 | 0 | 10884 | 103 | 11225 | 45 |
| 45967 | Caribou Flats, NT | 1188869 | 0.803903 | 48501 | 2.21679 | 559 | 0 | 391 | 0 | 10760 | 77 | 11371 | 42 |
| 45968 | Caribou Flats, NT | 828539 | 0.802416 | 34682 | 2.21652 | 610 | 0 | 384 | 0 | 10768 | 90 | 11151 | 48 |
| 45994 | Slate Islands, ON | 1170660 | 0.801806 | 48348 | 2.21754 | 412 | 1 | 281 | 0 | 8061 | 48 | 8404 | 32 |
| 48050 | South Slave, NT | 1179551 | 0.803121 | 48495 | 2.21728 | 534 | 0 | 371 | 0 | 10555 | 74 | 11241 | 38 |
| 48057 | South Slave, NT | 1144904 | 0.8023 | 47171 | 2.21662 | 569 | 0 | 382 | 0 | 10616 | 99 | 11151 | 53 |
| 48075 | South Slave, NT | 1156803 | 0.802571 | 47716 | 2.21711 | 560 | 0 | 385 | 0 | 10232 | 78 | 11035 | 46 |
| 48094 | South Slave, NT | 1149653 | 0.802727 | 47182 | 2.21795 | 582 | 0 | 396 | 0 | 10521 | 95 | 10917 | 54 |
| 48100 | Quintette, BC | 1135834 | 0.801004 | 46529 | 2.2182 | 546 | 0 | 366 | 0 | 10615 | 87 | 10889 | 42 |
| 48105 | Graham, BC | 1161224 | 0.801082 | 47638 | 2.21698 | 557 | 0 | 376 | 0 | 10123 | 81 | 10549 | 56 |
| 48111 | Graham, BC | 1134432 | 0.800771 | 46595 | 2.2174 | 536 | 0 | 350 | 0 | 10542 | 76 | 11202 | 60 |
| 48112 | Kennedy Siding, BC | 1185400 | 0.800698 | 48659 | 2.21808 | 524 | 0 | 348 | 0 | 10385 | 93 | 10906 | 56 |

**Supplementary Figures**

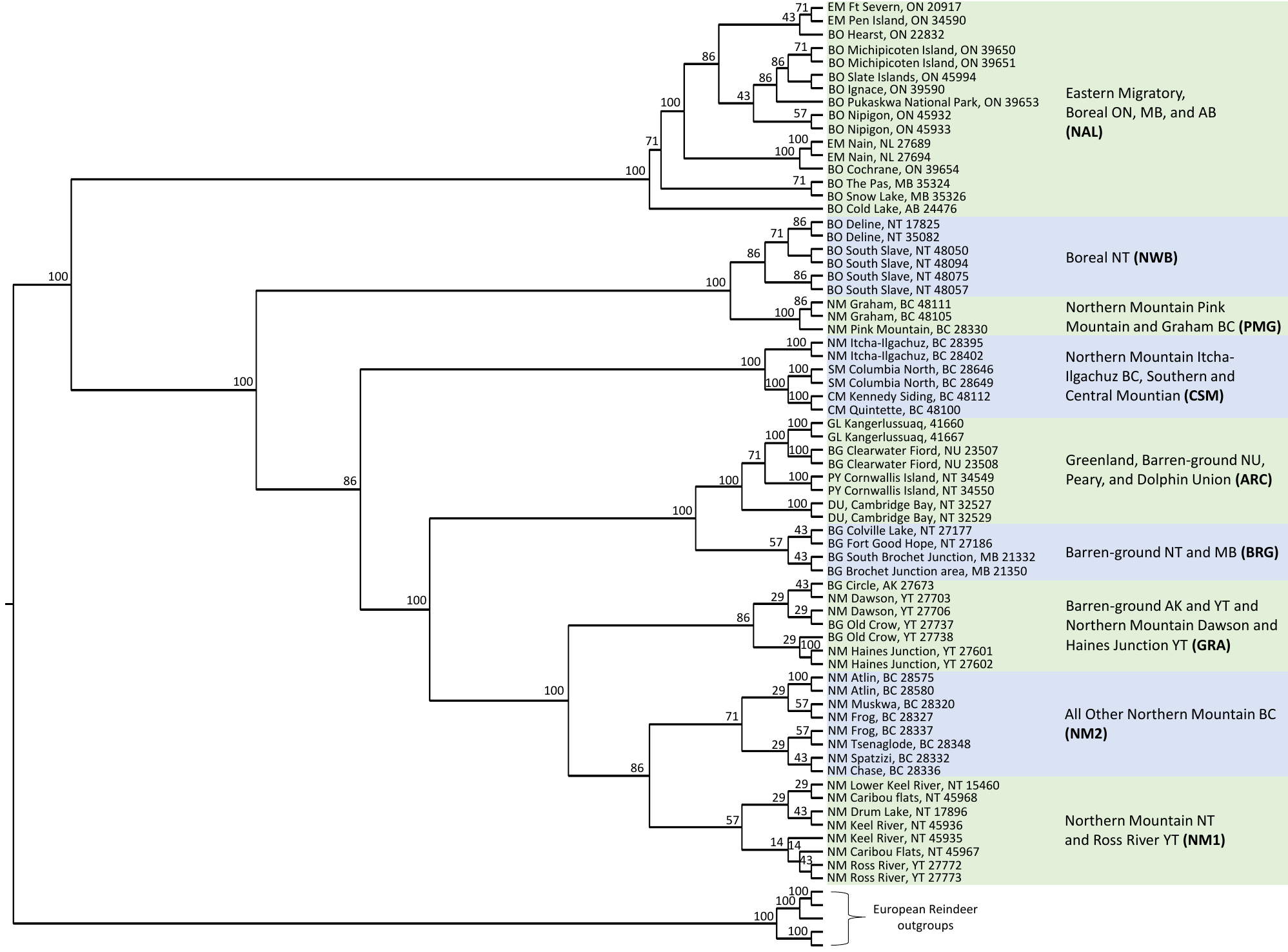

**Supplementary Figure 1**. Maximum likelihood phylogenomic reconstruction using the whole genome sequences of the 66 caribou and five reindeer genomes as outgroups. Bootstrap supports are shown on the nodes. Each individual is identified on the tree with their PCID number (Table S1) and their Designatable Unit; BG – barren-ground, BO – boreal, CM – central mountain, DU – Dolphin Union, EM – eastern migratory, GL – Greenland, NM – northern mountain, PY – Peary, and SM – southern mountain.

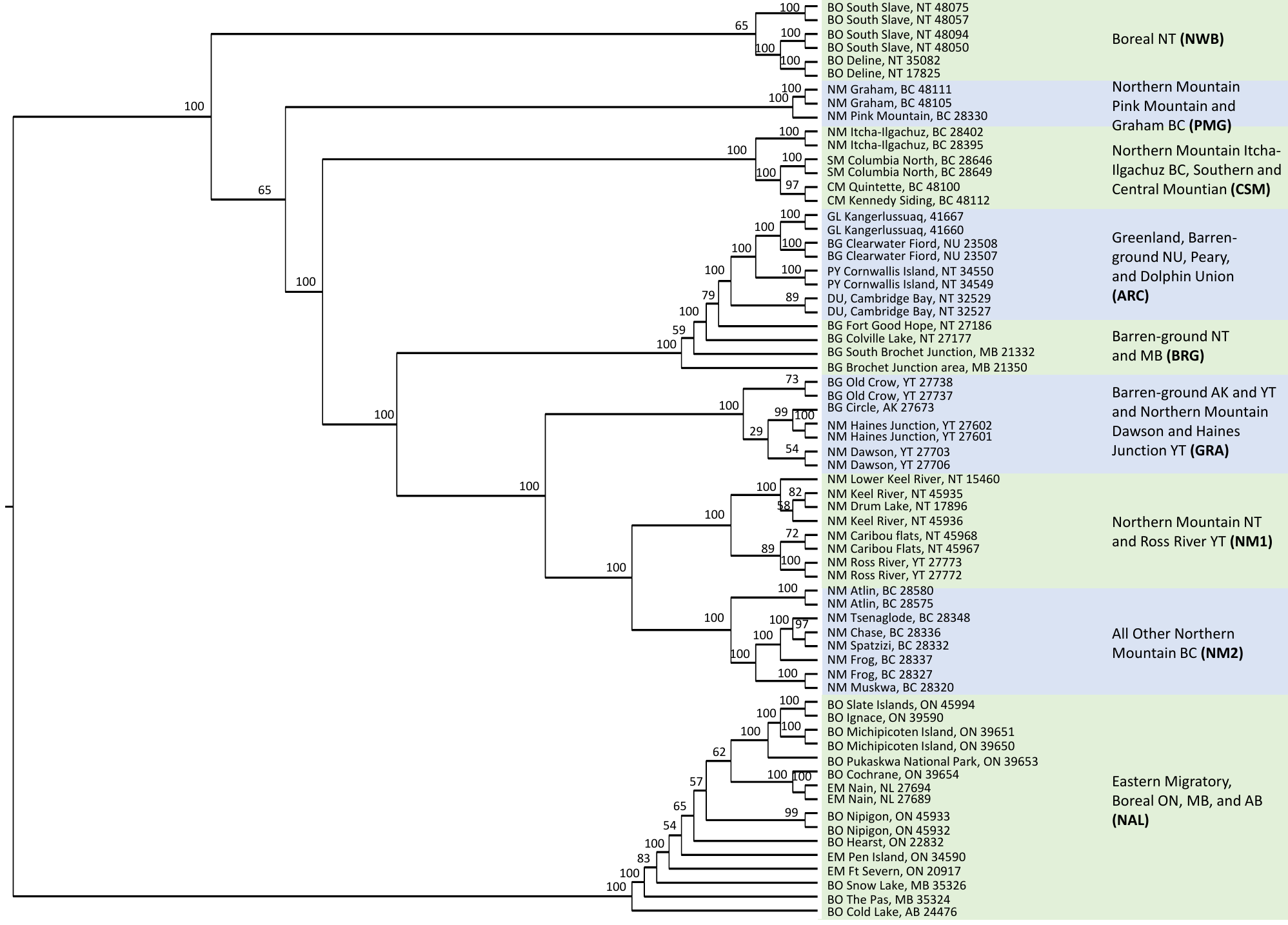

**Supplementary Figure 2.** SNP based phylogenomic reconstruction of the 66 caribou. Bootstrap supports are shown on the nodes. Each individual is identified on the tree with their PCID number (Table S1) and their Designatable Unit; BG – barren-ground, BO – boreal, CM – central mountain, DU – Dolphin Union, EM – eastern migratory, GL – Greenland, NM – northern mountain, PY – Peary, and SM – southern mountain

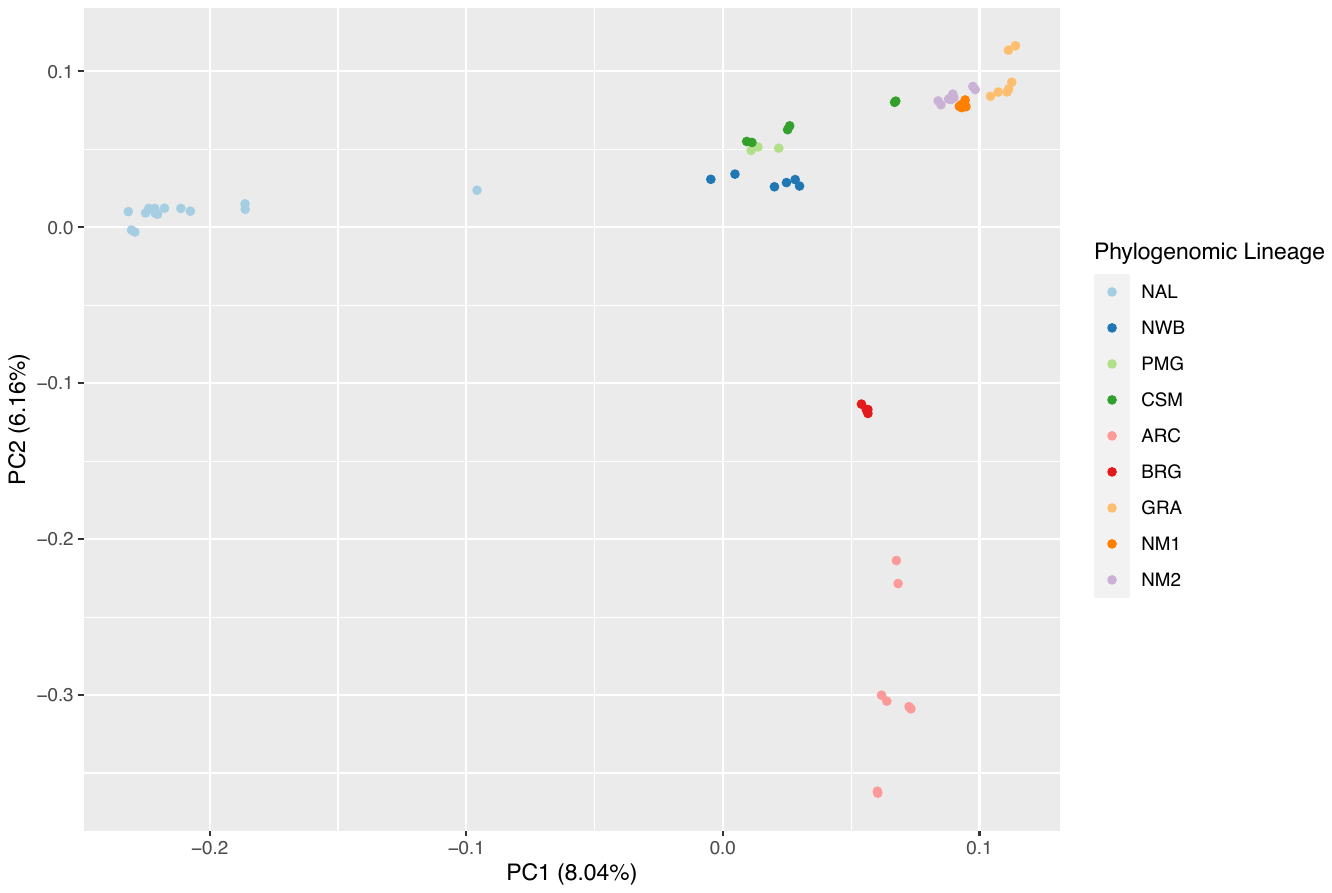

**Supplementary Figure 3.** PCA with all 66 caribou genomes, with the points coloured by the lineage within which it sits on the phylogenomic reconstruction.

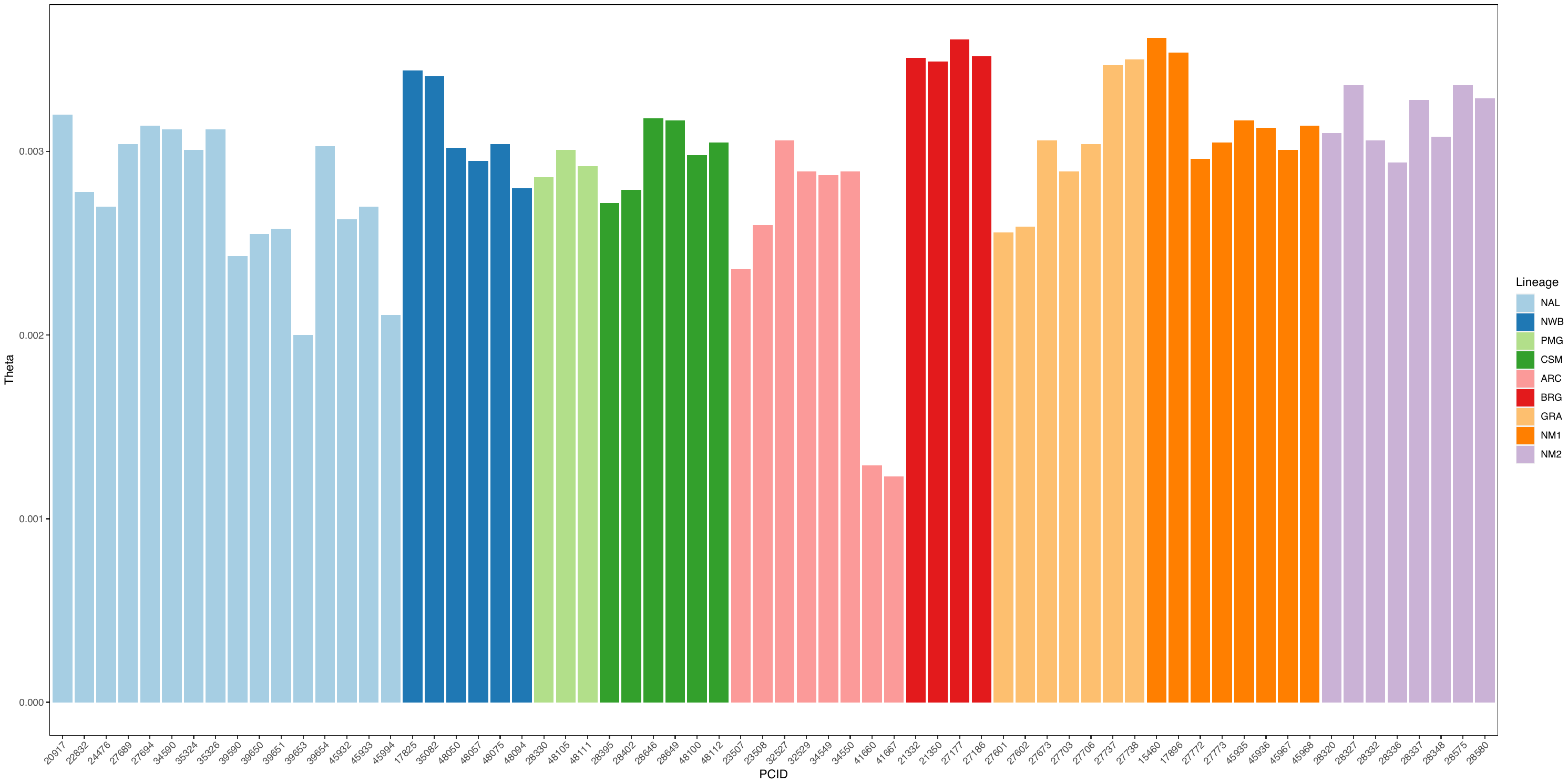

**Supplementary Figure 4.** Individual genetic diversity, θ, an approximation of heterozygosity.

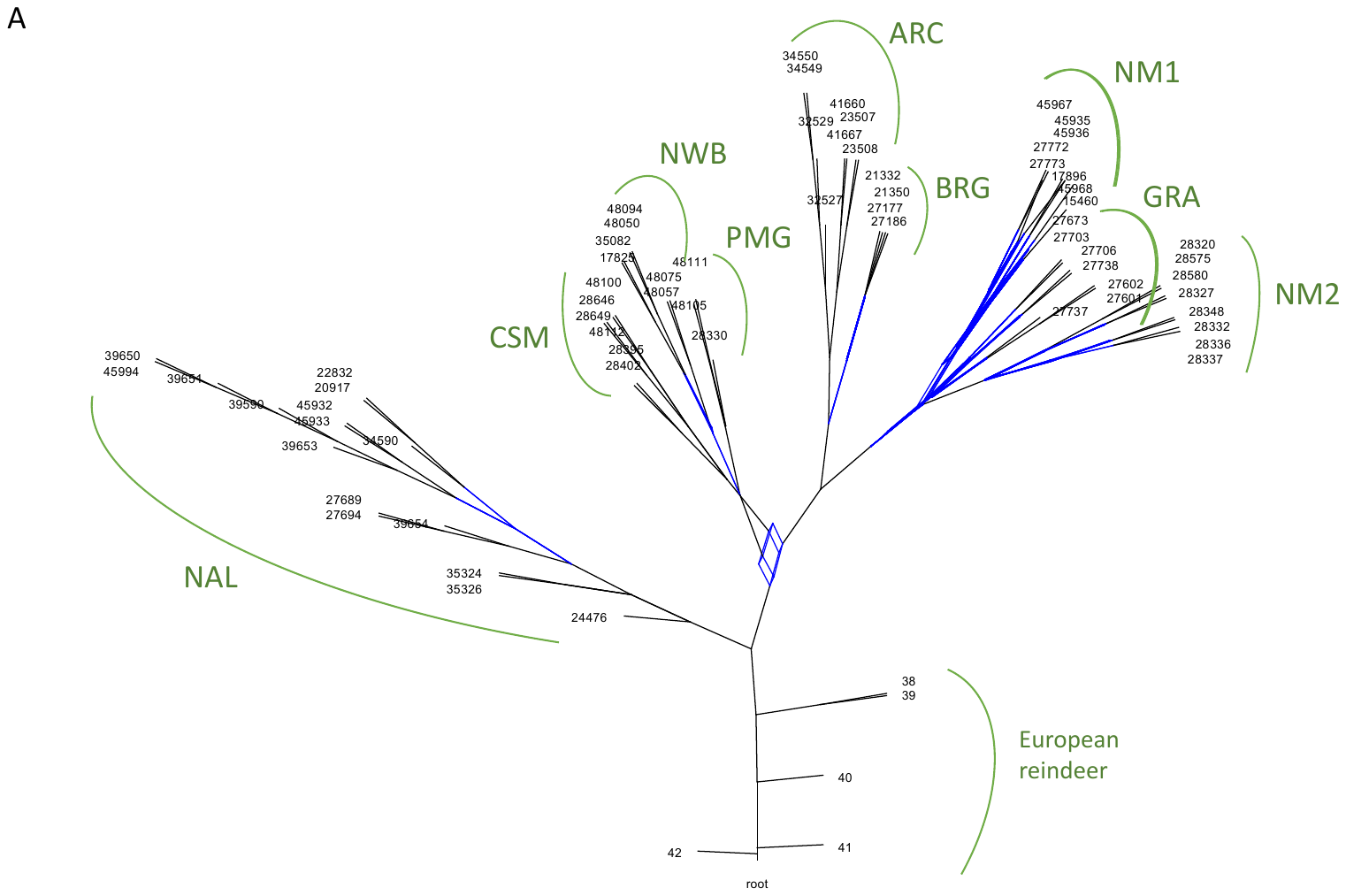

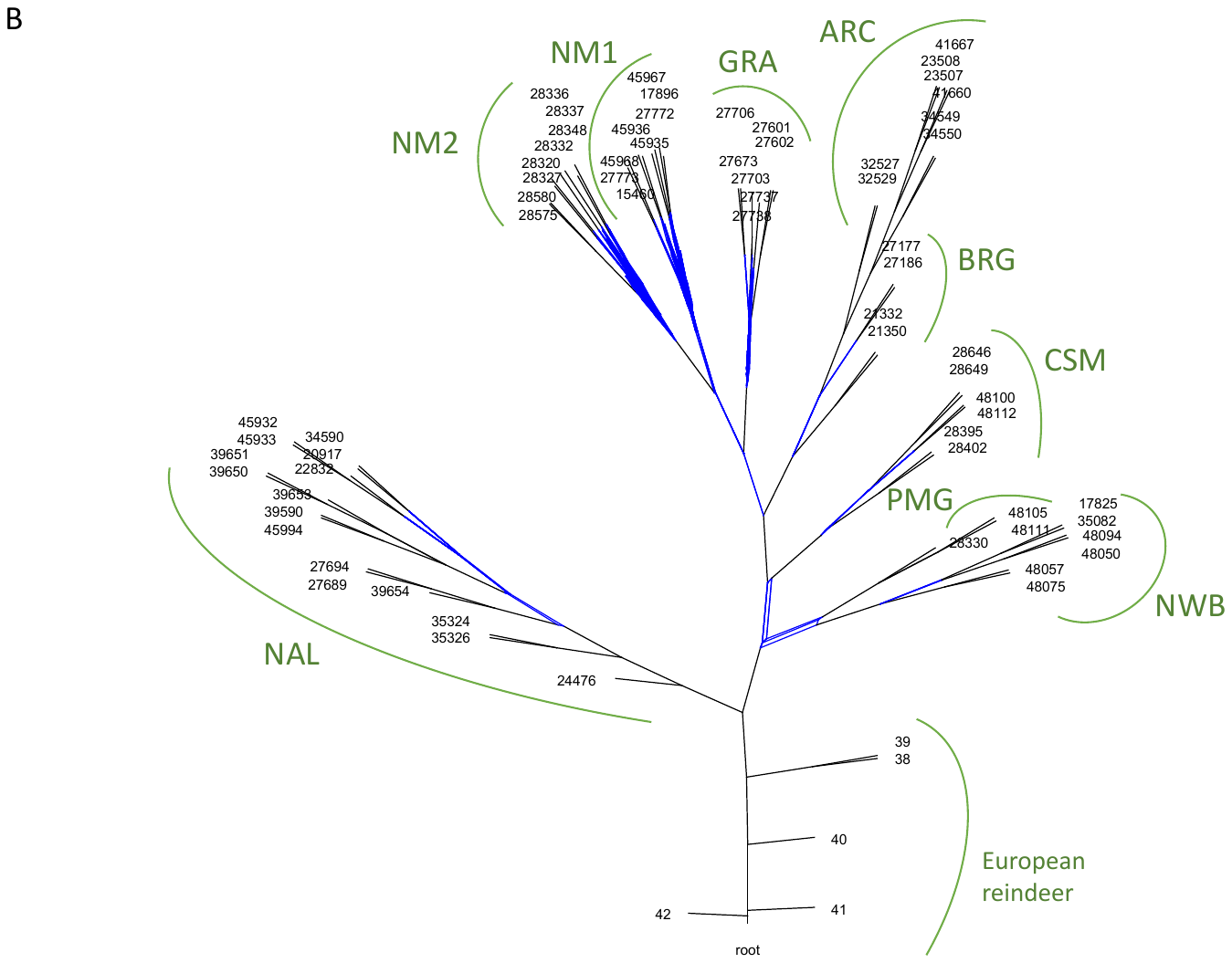

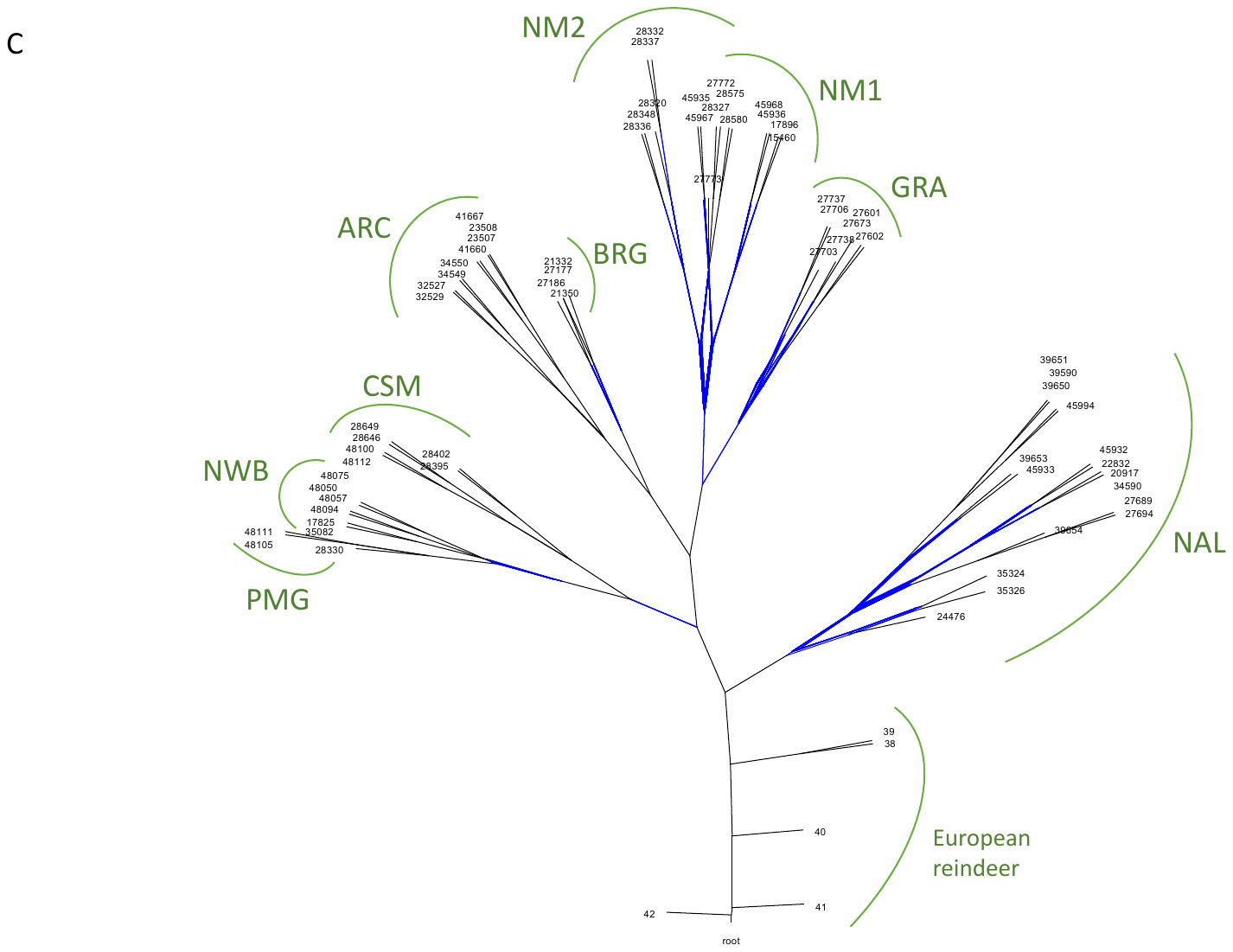

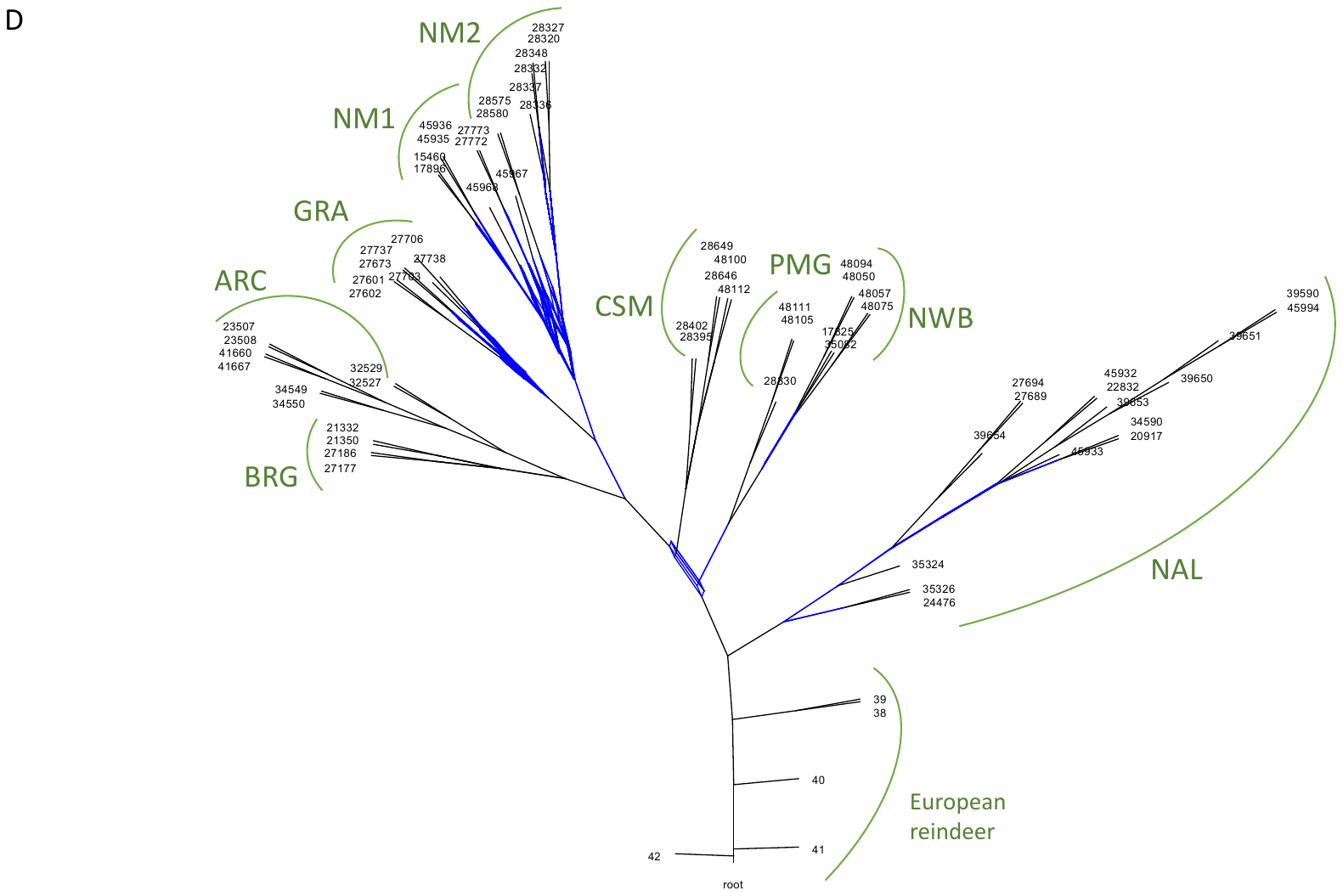

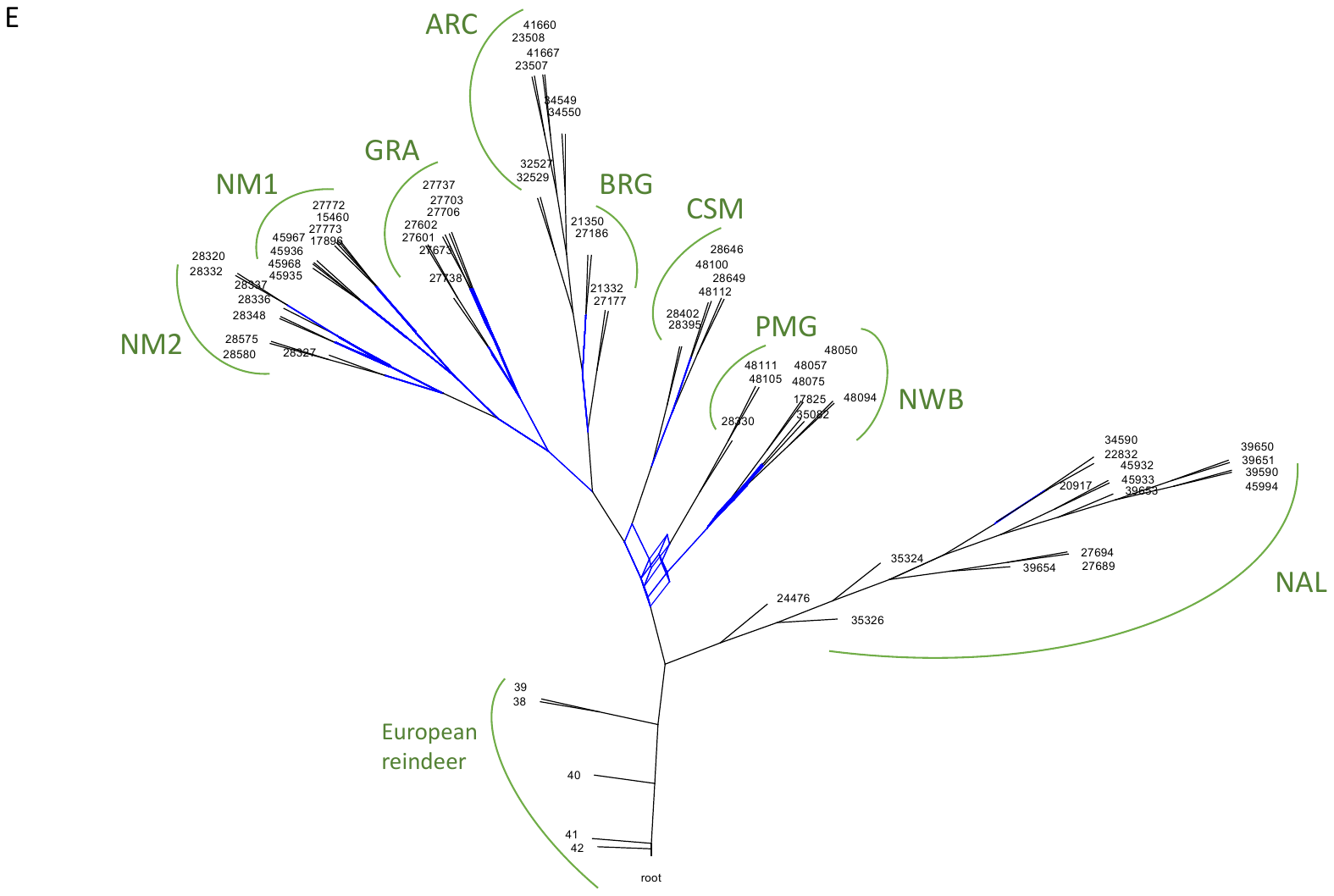

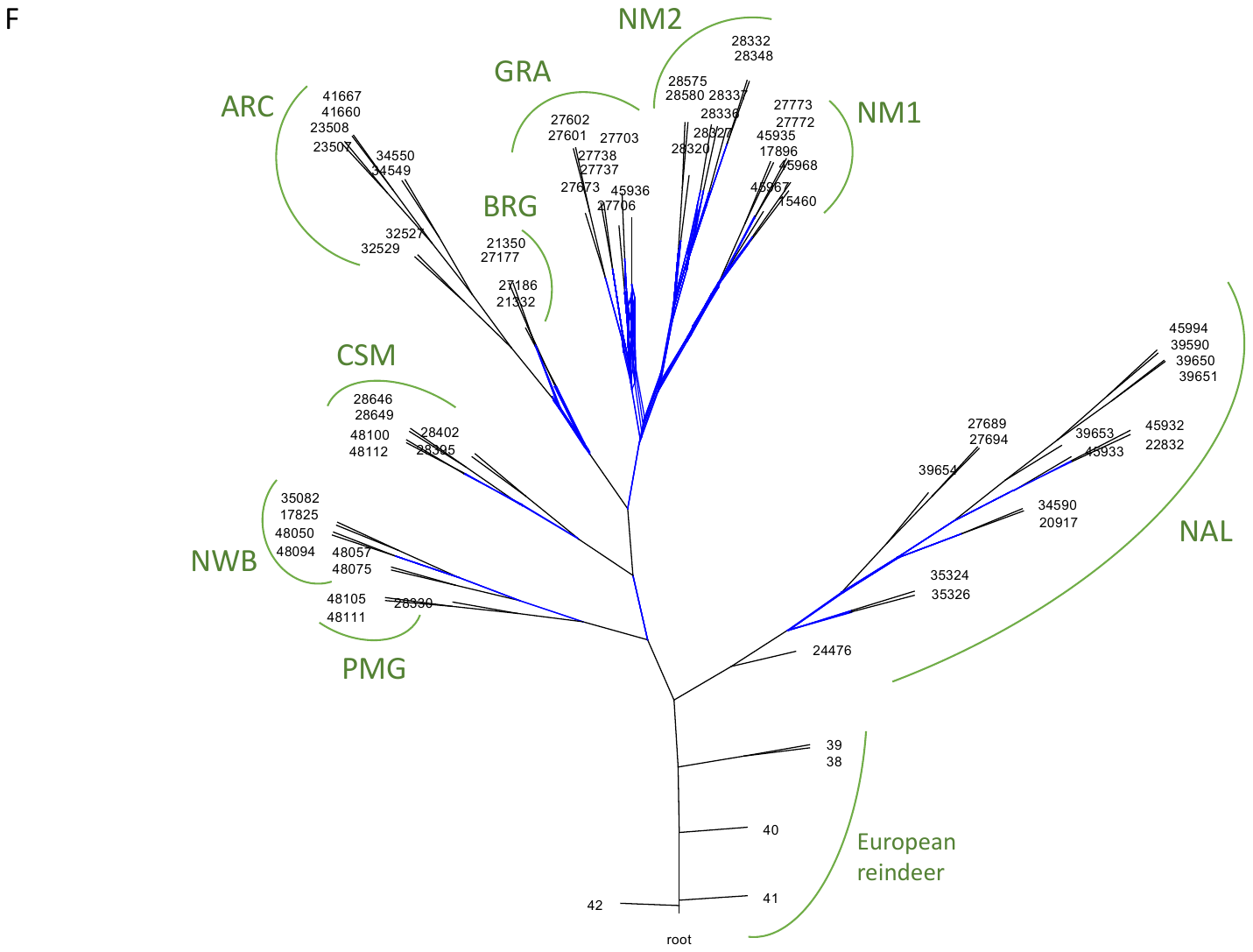

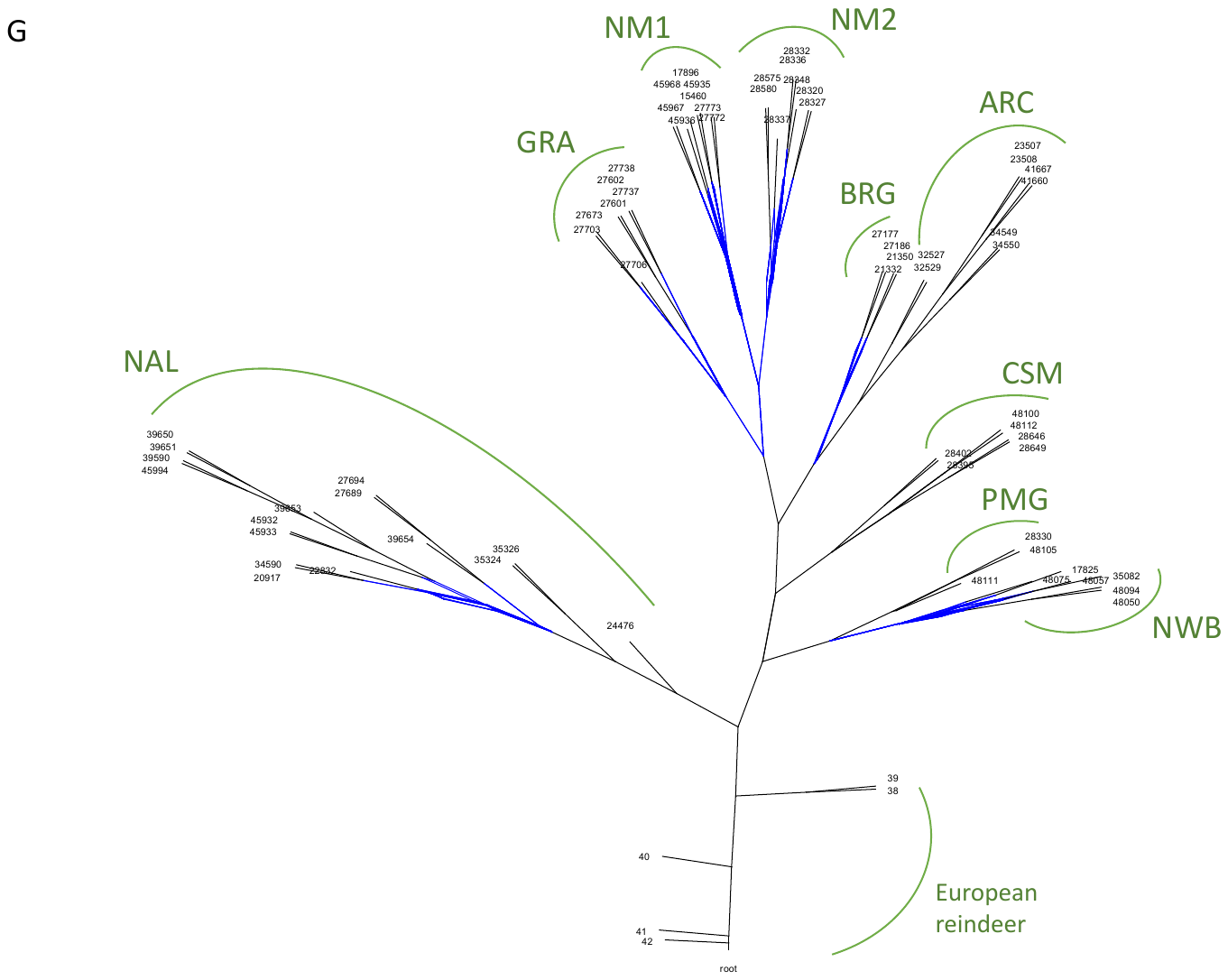

**Supplementary Figure 5.** Admixture graphs resulting from the phylogenomic reconstruction of 66 caribou using scaffolds 1-3 (**a**), 4-7 (**b**), 8-11 (**c**), 12-16 (**d**), 17-21 (**e**), 22-27 (**f**), and 28-35 (**g**). Blue lines represent phylogenetic uncertainty putatively resulting from gene flow. Green labels show the lineage to which the individuals belong.

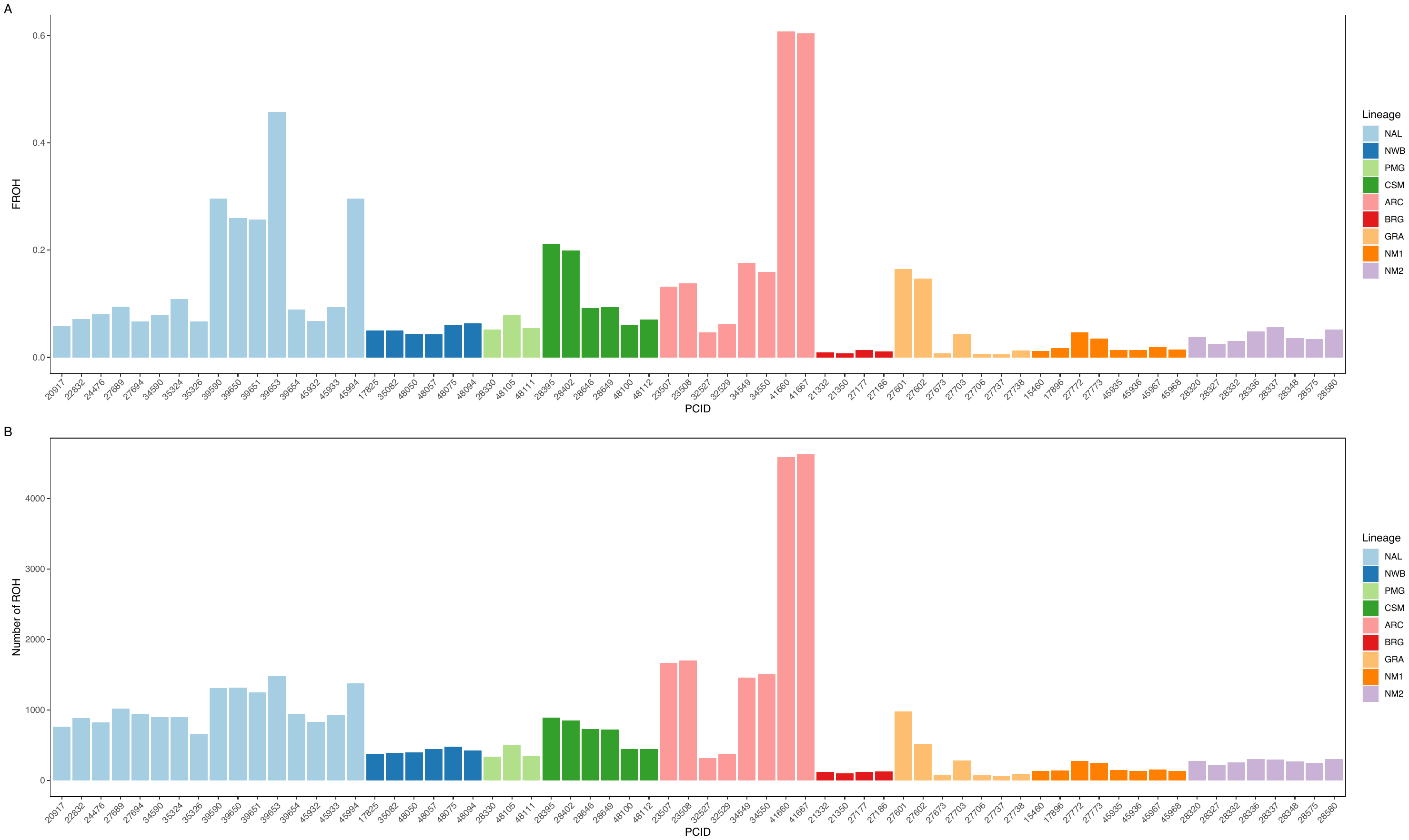

**Supplementary Figure 6**. The proportion of the genome in runs of homozygosity (FROH; **a**) and the number of runs of homozygosity for each individual (**b**).

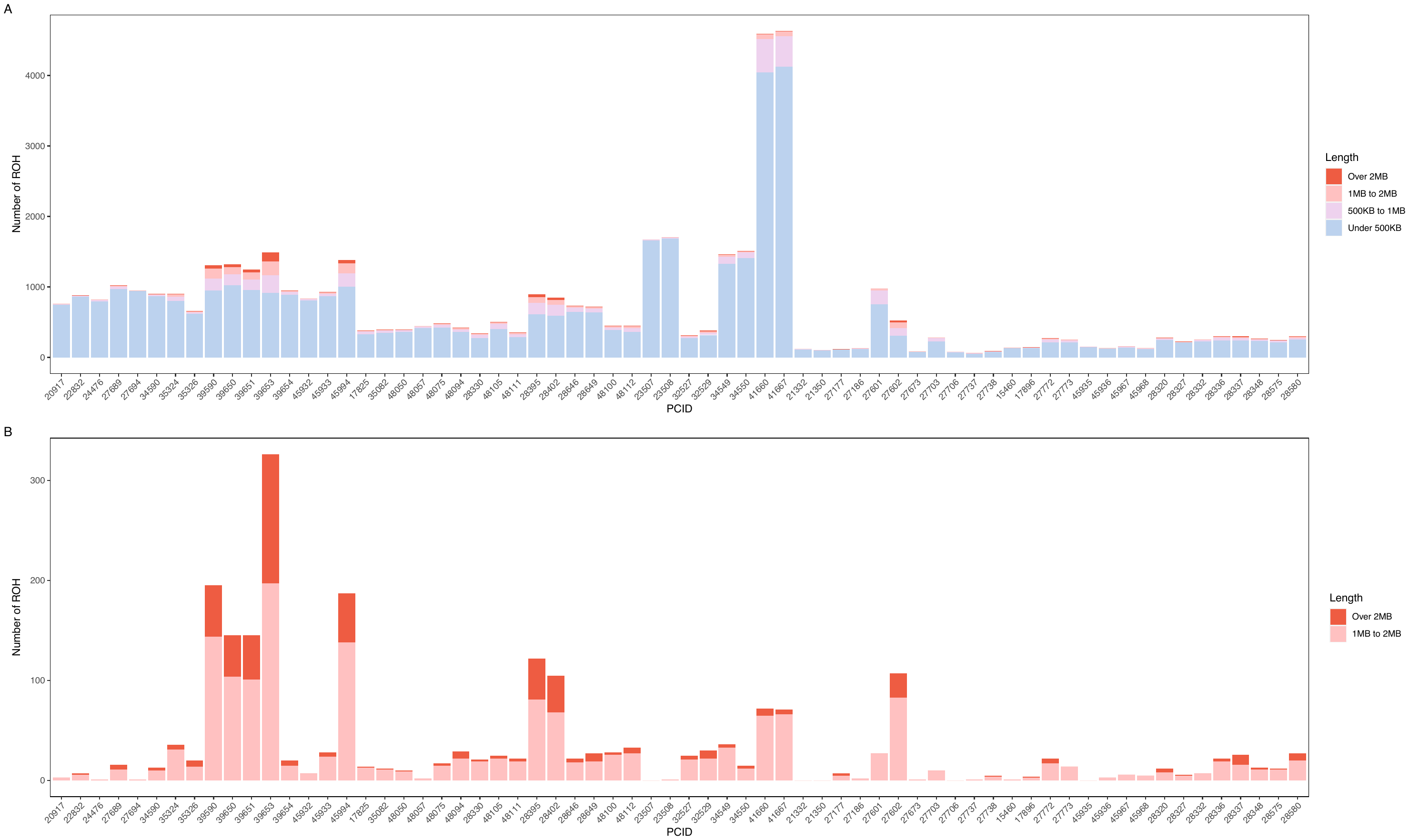

**Supplementary Figure 7**. The number of runs of homozygosity in each individual under different length classes (**a**) and the number of runs of homozygosity over 1 million base pairs (1MB) for each individual (**b**).

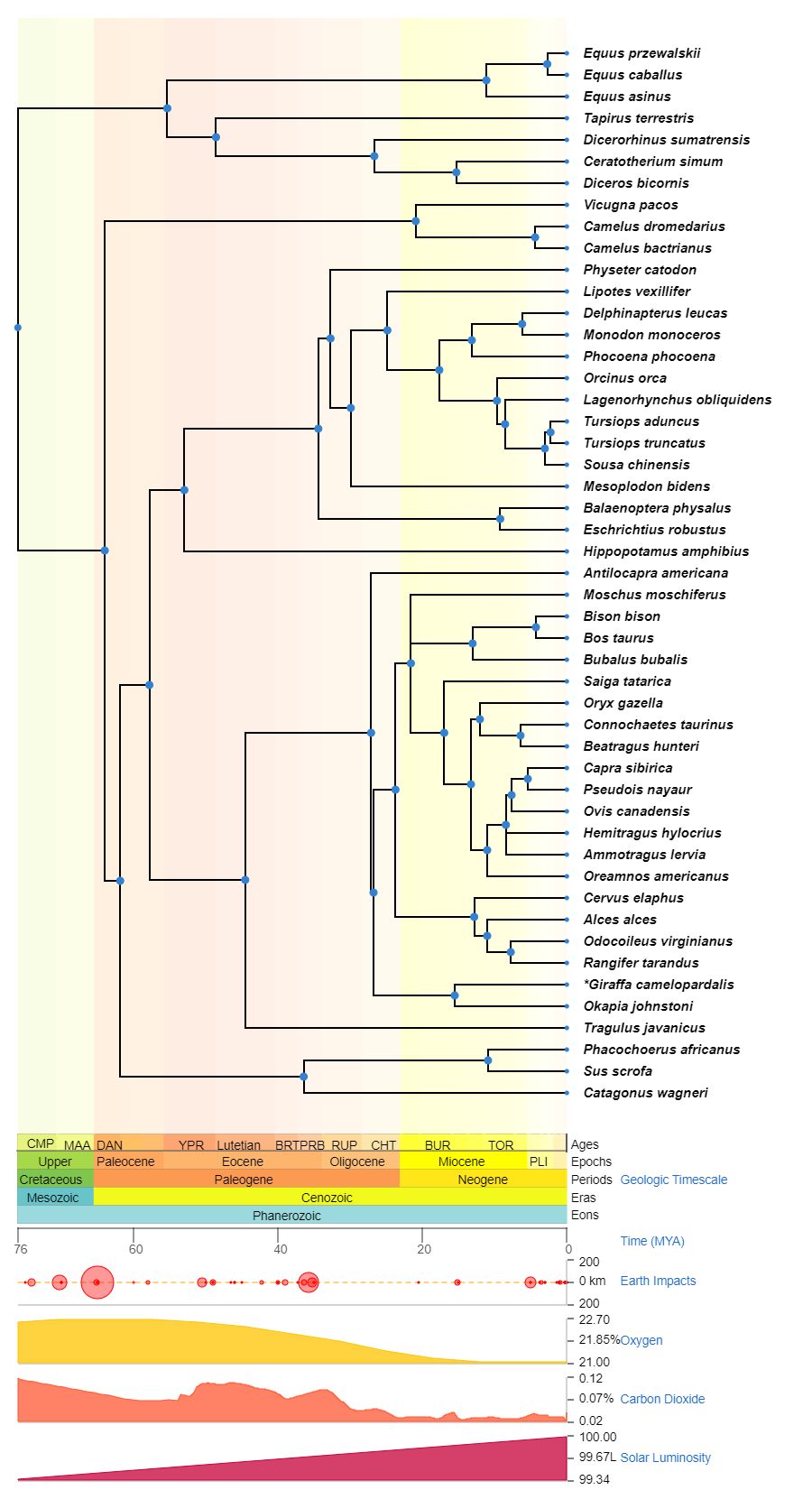

**Supplementary Figure 8**. TimeTree phylogenetic reconstruction of the mammal species used for the GERP analysis.
